# Molecular interplay between ComEC domains leads to efficient DNA translocation during natural transformation

**DOI:** 10.1101/2025.04.08.647572

**Authors:** Matthew J.M. Stedman, Sophie Deselaers, Sebastian A.G. Braus, Dianhong Wang, Maria Gregori Balaguer, Alvar D. Gossert, Manuela K. Hospenthal

## Abstract

Naturally competent bacteria can take up and incorporate free DNA from their environment using complex machinery that is endogenously encoded. This process is called natural transformation and is a key mechanism in the spread of antibiotic resistance amongst bacteria, including many human pathogens. All competent bacteria require ComEC to transport the transforming DNA across the cytoplasmic membrane. In addition to the transmembrane competence domain predicted to form the DNA channel, most ComEC orthologues additionally contain an oligonucleotide binding (OB) domain and β-lactamase-like domain. Here, we provide in-depth characterisation of the nuclease activity of the β-lactamase-like domain and the DNA binding activity of the OB domain, and present high-resolution structures of both domains. We show that the *in vitro* nuclease activity of the β-lactamase-like domain is enhanced when the OB domain is encoded on the same polypeptide chain. Additionally, we identify a pin loop within the β-lactamase-like domain responsible for melting the DNA duplex prior to cleavage of the non-translocating strand, and a DNA channel lined with aromatic residues that guide the uncleaved translocating strand through ComEC. On the basis of our biochemical, structural and functional characterisation, we provide a mechanistic model for how ComEC achieves the simultaneous tasks of DNA degradation and translocation, central to the natural transformation process.

## Introduction

Natural transformation is the uptake and subsequent genomic integration of exogenous DNA by naturally competent bacteria and, alongside transduction and conjugation, is a critical mechanism of horizontal gene transfer (HGT)(1, 2). HGT is a driver of bacterial evolution, with important consequences for the spread of antibiotic resistance and other pathogenicity traits(3, 4). Natural transformation can be distinguished from the other mechanisms of HGT, as the machinery required for DNA uptake, translocation and integration is entirely encoded, expressed and regulated by the competent cell(2, 5–7). To date, transformation has been directly observed in over 80 species, including both Gram-positive and Gram-negative organisms(7, 8), yet the true prevalence is likely much higher.

Although there are differences in the cell envelopes of Gram-positive and Gram-negative bacteria, the general mechanisms governing natural transformation are remarkably conserved. In most species, type IV pili or related structures mediate DNA capture and uptake into the periplasmic space(9). Most commonly the transforming DNA (tDNA) is linear and double stranded (dsDNA), although single stranded DNA (ssDNA) and circular plasmid DNA can also be taken up(10–12). Specialised pilus subunits capable of DNA binding have been identified in a number of species(13–20). In Gram-negative bacteria, DNA binding by the pilus and subsequent pilus retraction is thought to bring the DNA into proximity with the outer membrane (OM)(16). Pilus retraction is powered by the cytoplasmic ATPase PilT(21), yet the exact mechanisms of how the DNA is taken up across the OM and peptidoglycan layer into the periplasmic space, potentially through the secretin channel PilQ, remain to be defined. In the periplasmic space, DNA is bound by the DNA-binding protein ComEA, which ensures the unidirectional movement of DNA into the cell(22–26). ComEA is membrane-bound in Gram-positive bacteria and capable of forming homo-oligomers through an oligomerisation domain(26). This process is thought to condense the incoming DNA in the periplasm, thereby exerting a pulling force on the DNA. In the next step, a single strand of the transforming DNA is translocated across the cytoplasmic membrane, which is mediated by the putative channel protein ComEC, while the non-translocating strand is degraded(27–29). On the cytoplasmic side of the membrane, ComFC (ComF in most Gram-negative organisms)(30–32) and ComFA (a role potentially fulfilled by PriA in Gram-negative organisms)(33–37) act on the incoming ssDNA in ways that remain poorly understood. The ssDNA is protected from degradation by single-stranded DNA binding protein (Ssb) and DNA processing protein A (DprA)(38–40). DprA loads the DNA onto recombinase A (RecA), which can integrate the DNA into the host genome provided there is sufficient homology between the sequences(41). This process may further be assisted by the helicase ComM and the helicase-associated nuclease YraN(42–44).

ComEC is essential for natural transformation, with its deletion leading to total abrogation of transformation(24, 45–47). It is present in both Gram-positive and Gram-negative competent organisms and most ComEC orthologues contain three predicted domains: the transmembrane DNA channel or competence domain (Pfam database(48), PF03772), the oligonucleotide binding (OB) fold (PF13567; domain of unknown function 4131, DUF4131), and the β-lactamase-like domain (PF00753)(8, 49–51). Orthologues that lack either or both the OB fold and the β-lactamase-like domain have been identified in bioinformatic analysis(27), yet it remains to be shown whether all or some of the bacterial species that harbour these variants can undergo transformation. The competence domain appears to be universally conserved across competent species and is predicted to form the DNA channel critical for ComEC’s role of DNA transport across the cytoplasmic membrane(27–29). Likewise, removal of the OB fold from the *Bacillus subtilis* ComEC orthologue resulted in a strain unable to undergo transformation(28).

In 1995 it was reported that a *comEC* deletion strain of *B. subtilis* was able to bind more tDNA on its cell surface (via ComEA), but showed a dramatic effect on DNA internalisation(24). A few years later in 2001, *comEC* null mutations were shown to prevent degradation of the non-transforming strand(52). These observations led to the speculation that ComEC itself may harbour nuclease activity required for the degradation of the non-translocating strand. More recently, *in silico* predictions and subsequent confirmation of *in vitro* activity attributed this cryptic nuclease activity to the C-terminal β-lactamase-like domain of ComEC(29, 53). Importantly, mutations in the *B. subtilis* β-lactamase-like domain that affect catalytic activity were shown to affect transformation *in vivo*(53, 54). Interestingly, there are bacterial species capable of undergoing transformation that naturally lack the β-lactamase-like domain of ComEC. The *Streptococcus pneumoniae* ComEC orthologue likely contains an inactive β-lactamase-like domain, as it does not encode all of the predicted catalytic metal coordinating residues(29). In this organism, the degradation of the non-translocating strand is carried out by the membrane-bound EndA nuclease(55). The more distant *Helicobacter pylori* ComEC homologue neither contains the β-lactamase-like domain(45, 56), nor an EndA homologue, suggesting yet a further mechanism of strand degradation.

The β-lactamase-like domain of ComEC is a member of metallo-β-lactamase (MBL) superfamily(57). This superfamily of enzymes acts on a diverse set of substrates, including the β-lactam ring of the β-lactam class of antibiotics for which the name was coined. Within the MBLs, enzymes that hydrolyse the phosphoester bonds of a variety of substrates including nucleic acids and nucleotides, phospholipids and phosphonates make up the most widespread functional group of MBLs(58). The majority of nucleic acid processing MBLs are binuclear Zn^2+^-dependent enzymes, although other metal ions such as manganese, iron or magnesium also occur in MBLs(58–60). In 2021, Silale and colleagues showed that the nuclease activity of the β-lactamase-like domain of the thermophilic Gram-positive organism *Moorella thermoacetica* DSM 521 is dependent on Mn^2+^ coordination(53). The general catalytic mechanism of nuclease MBLs, such as RNAse J and RNAse Z, relies on deprotonation of an active site water molecule by an Asp general base. The metal ions coordinate the resulting hydroxide ion, which then acts as the nucleophile attacking the scissile phosphate of the substrate(61–65).

In this study, we provide an in-depth characterisation of the nuclease activity of the β-lactamase-like domain and show that it functions as both an endo- and exonuclease, the latter with exclusive selectivity for the 5′ terminus. We identify a pin loop in the β-lactamase-like domain responsible for the separation of double-stranded DNA prior to the hydrolysis of the strand harbouring the 5′ terminus. Subsequently, a series of aromatic residues that line the ssDNA channel guide the uncleaved translocating DNA strand through the competence domain. While the DNA binding affinity of the β-lactamase-like domain is undetectably weak *in vitro*, the OB fold tightly interacts with DNA. We show that when both domains are encoded on the same polypeptide chain, the OB fold increases the local concentration of the DNA substrate in proximity with the β-lactamase-like domain’s active site, thus enhancing its nuclease activity markedly. Together with *in vivo* functional assays and the structural characterisation of both the β-lactamase-like domain and the OB domain, we propose a mechanistic model of how ComEC functions to cleave the non-translocating strand, while the other is successfully translocated across the cytoplasmic membrane.

## Results

### The β-lactamase-like domain of ComEC degrades DNA through a two-metal-ion catalytic mechanism

Most ComEC orthologues are comprised of three domains: an OB fold encoded near the N-terminus, a central competence domain consisting of a bundle of transmembrane helices forming a channel and a C-terminal β-lactamase-like domain (**Fig. 1a**). In order to learn more about the mechanism of the β-lactamase-like domain, we sought to determine its three-dimensional structure. Previous work led to the identification of the *M. thermoacetica* β-lactamase-like domain orthologue as a well-expressed and soluble construct suitable for enzymatic characterisation(53). In our hands, the orthologue from *M. glycerini* (hereafter BLACT_Mg_), a close relative of *M. thermoacetica*, also yielded soluble and stable preparations, and proved to be amenable to structure determination by X-ray crystallography. We determined the 1.8 Å crystal structure of BLACT_Mg_, consisting of the characteristic αβ/βα fold that defines the MBL superfamily(57) (**Fig. 1b, Table S1**). The active site is positioned at one end of a wide shallow groove between the sandwiched central β-sheets. Our structure contains clear density for a phosphate ion in the active site, but lacks coordinated metal ions and thus represents an inactive state. A DALI(66) search revealed that the closest structural homologues of BLACT_Mg_ is teichoic acid phosphorylcholine esterase (Pce)(67) from *S. pneumoniae* and the closest nuclease is RNAse J from *Streptomyces coelicolor*(61) (**Fig. S1**). Like most MBLs with RNA or DNA substrates, RNAse J utilises two metal ions for catalysis(58–60) and its active site is situated at the interface between the hydrolytic β-lactamase domain and an auxiliary β-CASP domain. Indeed, nucleic acid specific members of the MBL superfamily often contain additional domains involved in substrate recruitment, such as the tRNAse Z exosite for tRNA binding(68), β-CASP(60) or KH(69, 70) domains for DNA or RNA binding. These domains can be inserted within the MBL fold, or occur N- or C-terminally to it(58). In the context of full-length ComEC, the OB fold may serve as the DNA binding domain that is lacking within the β-lactamase-like domain. While our structure of BLACT_Mg_ does not contain the β-CASP domain, it does display common sequence motifs found in other MBL enzymes(71), such as the characteristic H-X-H-X-D-H (where X is any residue) motif which consists of H598-X-D600-X-D602-H603 in BLACT_Mg_.

**Fig. 1:**
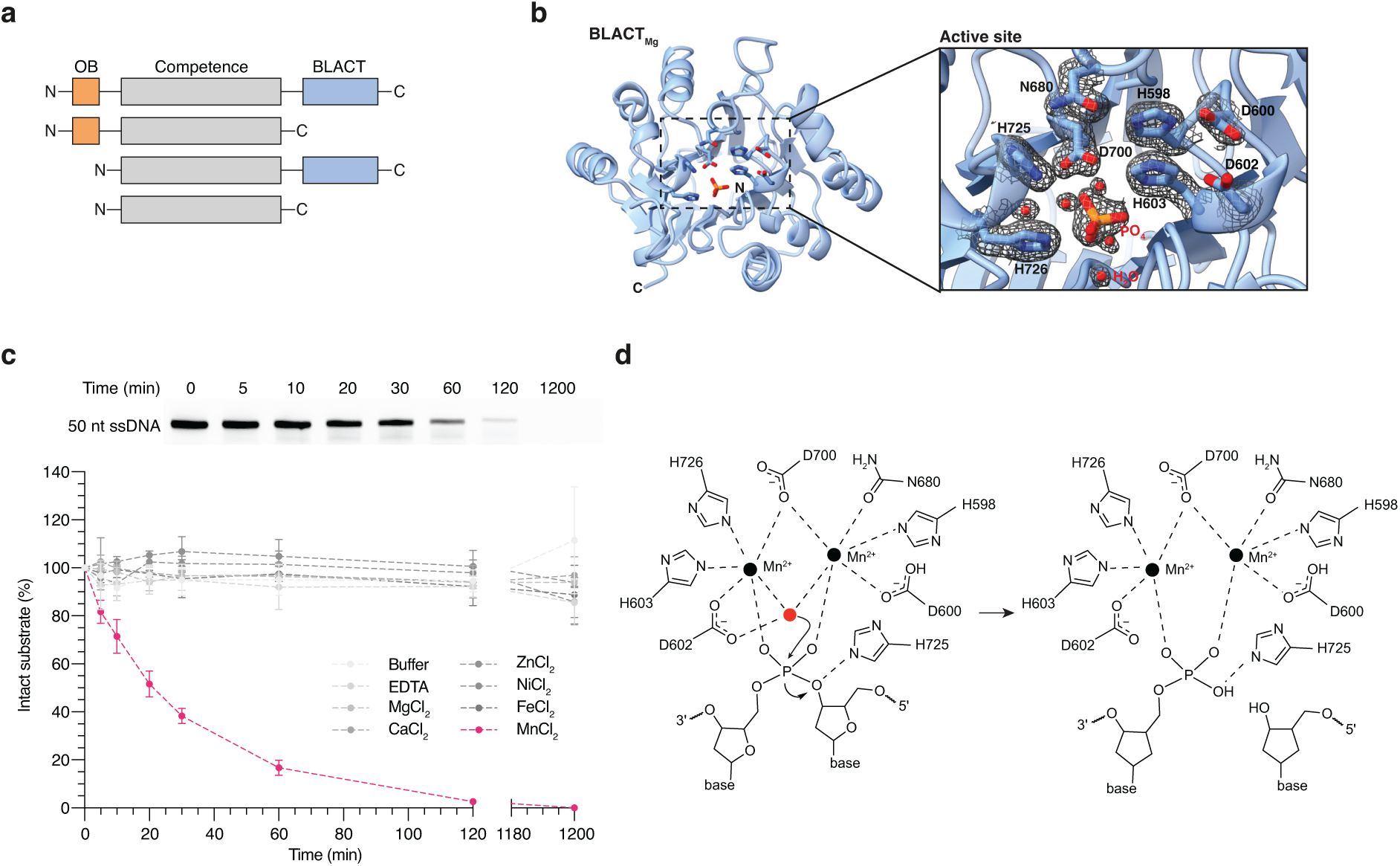
The structure of BLACT_Mg_ reveals the catalytic mechanism of ComEC’s β-lactamase-like domain. **a** Schematic illustrating the domain organisation of ComEC. Most ComEC orthologues are composed of three domains: the OB fold, the competence/channel domain and the β-lactamase-like domain, although versions lacking either or both the OB fold or β-lactamase-like domain also exist. **b** The crystal structure of BLACT_Mg_ displayed in ribbon representation with active site residues and the bound phosphate ion shown in stick representation. N- and C-termini are indicated on the structure. The inset shows a zoomed in view of the active site with ordered water molecules and the bound phosphate ion. Active site residues and the phosphate ion are shown in stick representation. The electron density around active site residues, water molecules and the phosphate ion is shown as a black mesh with the refined 2mF_O_-DF_C_ map contoured at 1 σ. **c** Time-course nuclease activity assay monitoring the degradation of ssDNA by BLACT_Mg_ in the presence of 5 mM of various metal ion cofactors. The image of the gel (top) was used to measure the fluorescence intensity of each band, from which the percentage of intact substrate remaining at each time point was plotted (bottom). Nuclease assays were conducted with 1 µM wild-type BLACT_Mg_ and 10 µM ssDNA substrates, 50 nt in length. DNA substrates were fluorescently labelled with fluorescein (FAM) on the first base (thymine, T) at the 5’ terminus. Error bars represent the standard error from three technical replicates. **d** Schematic showing the proposed catalytic mechanism of BLACT_Mg_. Deprotonation of a coordinated water molecule (red) by D602 functioning as a general base forms a hydroxide ion bridging the two active site metals. This ion initiates a nucleophilic attack on the scissile bond resulting in new 5’ phosphate and 3’ OH groups through nucleophilic substitution.

In a previous study it was observed that the β-lactamase-like domain from *M. thermoacetica* is Mn^2+^-dependent(53). We were unable to produce crystals in the presence of supplemented Mn^2+^, and, accordingly, there is no density that would correspond to the coordinated metal ions. In order to test whether BLACT_Mg_ is also a Mn^2+^-dependent nuclease, we performed a gel-based time-course nuclease activity assay using fluorescently labelled 50 nucleotide (nt) ssDNA as the substrate in the presence of various potential metal cofactors, followed by fluorescence intensity measurement of the band corresponding to the original intact substrate (**Fig. 1c**). The fluorescein (FAM) moiety was attached on the first base (thymine, T) at the 5’ terminus. These assays indeed confirmed that, like its close relative from *M. thermoacetica*, BLACT_Mg_ is also an Mn^2+^-dependent nuclease. The gradual decrease in band intensity without a change in its electrophoretic mobility also suggests that no or very little 3’-to-5’ exonuclease activity is taking place in this reaction. To definitively determine the coordination number of BLACT_Mg_, we performed mass spectrometry on our purified BLACT_Mg_ sample, with and without supplementary Mn^2+^ in the solution (**Fig. S2**). BLACT_Mg_ contains no bound Mn^2+^ without specific supplementation of the metal (as captured by our crystal structure), whereas the majority of the molecules in our sample are coordinated with two Mn^2+^ ions when BLACT_Mg_ was incubated with 5 mM MnCl_2_. The absence of coordinated Mn^2+^ ions prevents the active site from adopting a fully catalytically competent conformation. In order to generate a general model for catalysis and to better understand the active site residue rearrangements necessary, we compared the active site of BLACT_Mg_ with structures of other MBL family enzymes where two metal cofactors were coordinated in the crystal structure (**Fig. 1d**, **Fig. S3**). We propose that D602, H603 and D600 rotate into the catalytic centre to coordinate metal ions. This would place the substrate in proximity with H725, the metal ions, and an active site water, allowing for catalysis to proceed. In this conformation, deprotonation of the active site water by the general base (D602) could proceed resulting in the formation of a hydroxide ion. This hydroxide bridges the two Mn^2+^ ions in the active site prior to nucleophilic attack on the 5′ side of the scissile phosphate. The nucleophilic substitution generates the 5′ phosphate and 3′ OH products. In general, the residues in and around the active site are highly conserved (**Fig. S4**).

In order to test the effect of mutation of important residues on nuclease activity, we performed further nuclease activity assays comparing wild-type BLACT_Mg_, a control mutation (D541A) (highly conserved but not in the active site), several residues involved in Mn^2+^-coordination (H598A, D600A, D602A, H603A, D700A) and putative DNA binding residues (H725A, S728A) (**Fig. S5, S6**). Mutation of most residues involved in metal coordination render BLACT_Mg_ virtually inactive in our nuclease activity assay, with the only exception being D600A that showed a ∼5 fold reduction in the apparent initial rate compared to wild-type. Similarly, substitution of the putative DNA binding residue S728 to alanine resulted in a ∼3 fold reduction in apparent initial rate, whereas the control mutation D541A showed similar activity compared to the wild-type. We also tested the effect of these mutations on transformation efficiency *in vivo*, where the analogous residues in *Legionella pneumophila* Lp02 were targeted (**Fig. S5**). Interestingly, all residues that showed reduced nuclease activity *in vitro*, except for S662A (corresponding to S728 in *M. glycerini*), showed complete abrogation of transformability. Similar mutations of catalytic β-lactamase-like domain residues in *B. subtilis* also resulted in reduced transformation efficiencies(53, 54), albeit to a lesser extent. Taken together, BLACT_Mg_ coordinates two Mn^2+^ in its catalytic centre in order to function as a DNA nuclease. When critical residues involved in the catalytic mechanism are substituted to alanine, BLACT_Mg_ is mostly rendered inactive both *in vitro* and *in vivo*, resulting in a severe transformation phenotype.

### BLACT_Mg_ is an endonuclease and 5′-to-3′ exonuclease *in vitro*

At present it is unclear whether the β-lactamase-like domain of ComEC encounters ssDNA or dsDNA *in vivo*. Even when dsDNA is taken up into the periplasmic space, it is possible that the translocating and non-translocating strands are separated prior to encountering the nuclease domain, which then engages with a single stranded substrate. Furthermore, we reasoned that the physiological DNA substrate of the β-lactamase-like domain *in vivo* contains a phosphate group at the 5′ end. In order to test whether BLACT_Mg_ shows an intrinsic preference for ssDNA or dsDNA and whether the presence of a hydroxyl or phosphate group at the 5’ end of the DNA substrate influences the rate of DNA degradation, we performed another time-course nuclease activity assay (**Fig. 2a**). In order to maintain the total number of cleavable phosphodiester bond the same between ssDNA and dsDNA substrates, we used 10 µM ssDNA and 5 µM dsDNA in this experiment. These data showed that BLACT_Mg_ has a clear preference for ssDNA and that the rate of degradation is slightly increased when ssDNA is modified with a 5’ phosphate group. This is consistent with the observation that the closest structural homologues of BLACT_Mg_ are enzymes that degrade single-stranded nucleic acid substrates and that the most likely physiological substrate is also the best substrate.

**Fig. 2:**
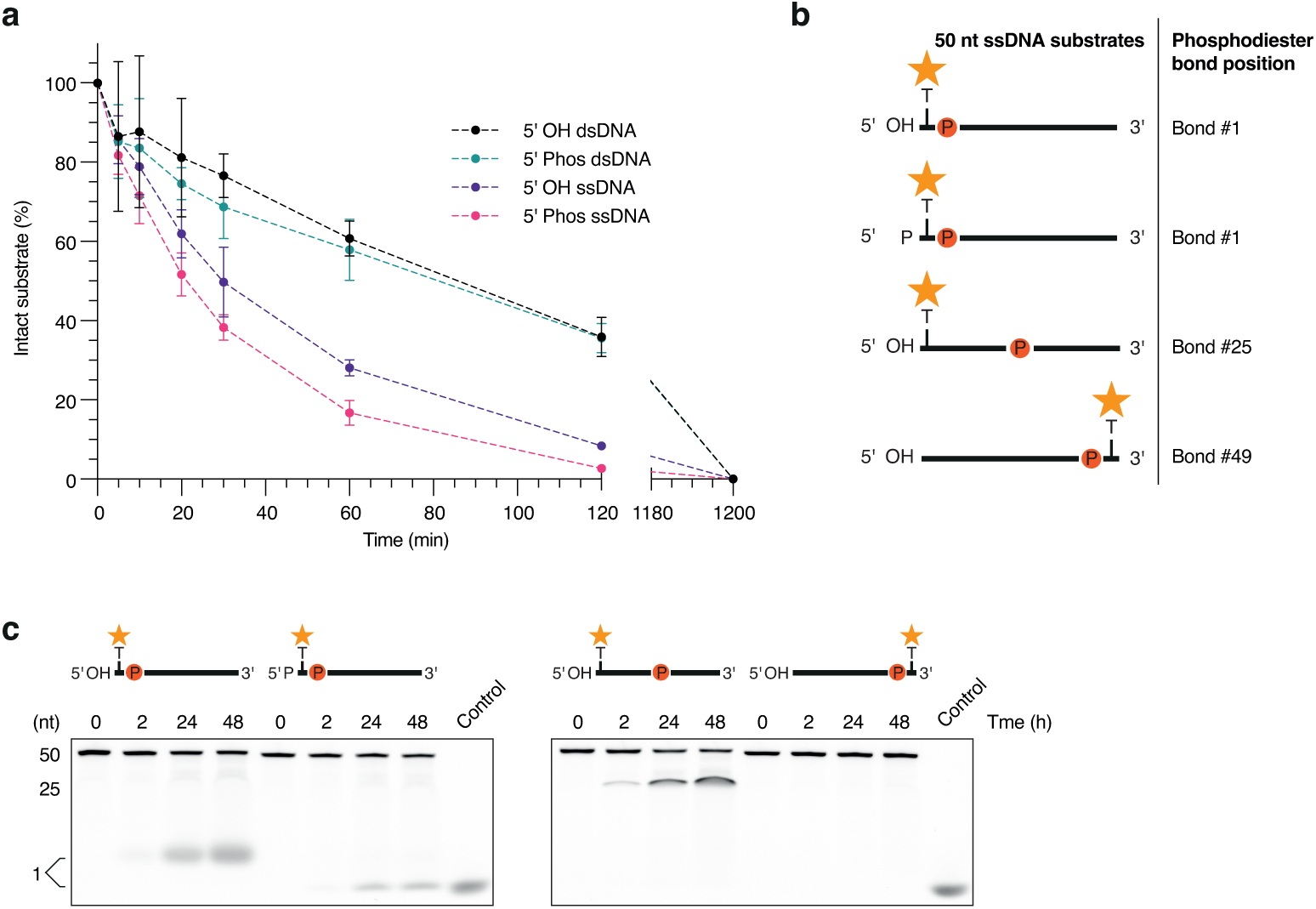
BLACT_Mg_ displays endonuclease and 5′-to-3′ exonuclease *in vitro*. **a** Time-course nuclease activity assay comparing the degradation of 5’ OH or phosphate-modified ssDNA or dsDNA. These experiments were performed using 10 µM ssDNA (50 nt) or 5 µM dsDNA (50 bp) in order to keep the total number of cleavable phosphodiester bonds identical. The mode of FAM-labelling was identical to the experiments in Figure 1. Error bars represent the standard error from three technical replicates. **b** Schematic illustrating the DNA substrates utilised in the experiment. All 50 nt ssDNA substrates are FAM-labelled (orange star), either at the first or last base (thymine, T) reading in the 5′ to 3′ direction. Each substrate carries a single phosphodiester bond marked by the letter P inside a red circle, while all other backbone sugar moieties are linked by nuclease-resistant phosphorothioate (PTO) bonds. **c** BLACT_Mg_-mediated cleavage of the four substrates depicted in (**a**) monitored over a 48 h time course resulting in the appearance of distinct cleavage products. Control, fluorescein-labelled dUTP (fluorescein-12-dUTP).

Our nuclease activity assay monitors the disappearance of the original intact substrate provided, namely FAM-labelled 50 nt ssDNA. Since the FAM moiety is attached to the first base at the 5’ end of the DNA molecule, we would observe a decrease in band intensity if BLACT_Mg_ degrades the substrate either with an endonucleolytic or 5’-to-3’ exonucleolytic cleavage mode. In contrast, if the substrate were degraded in a 3’-to-5’ exonucleolytic mode, we would see no change in band intensity, but rather a gradual increase in its electrophoretic mobility (gradual downward shift). As alluded to above, we do not observe such a change in electrophoretic mobility, suggesting that BLACT_Mg_ uses an endo- and/or 5’-to-3’ exonucleolytic mode. The slight preference for DNA substrates modified with a phosphate group at the 5’ terminus suggests that at least some exo-activity occurs, as endo cleavage events should not be affected by the terminus modification potentially some distance removed. In order to learn more about the cleavage mode of BLACT_Mg_, we designed several distinct ssDNA substrates consisting of nuclease-resistant phosphorothioate (PTO) bonds and a single cleavable phosphodiester bond and performed a nuclease assay (**Fig. 2b, c**). The cleavable bond was positioned either at the 5’ or 3’ terminus, or in the middle of the 50 nt ssDNA, and the FAM label was attached such that we could monitor the appearance of a single nucleotide product (**Fig. 2b**). These experiments showed that BLACT_Mg_ is able to cleave the first nucleotide from the 5′ end exonucleolytically, regardless of whether the terminus is modified by a phosphate or hydroxyl group. In contrast, there is no exonucleolytic cleavage occurring from the 3′ terminus, even after 48 h. Lastly, the enzyme is also clearly capable of cleaving endonucleolytically, which was further confirmed by the ability of BLACT_Mg_ to degrade plasmid DNA (**Fig. S7**), as shown previously(53). In summary, *in vitro* the BLACT_Mg_ functions as a 5′-to-3′ exonuclease and an endonuclease.

### The OB fold of ComEC binds to DNA with high affinity through an electrostatically driven interaction

In addition to the β-lactamase-like domain and the competence/channel domain, the majority of ComEC orthologues were found to also contain an OB fold encoded near the N-terminus(27) (**Fig. 1a**). Structural predictions suggest that the OB fold of Gram-positive organisms like *M. glycerini* is located on the extracellular side of the cytoplasmic membrane(28), and in Gram-negative organisms like *L. pneumophila* it is located in the periplasm (**Fig. S8**). In addition, the OB fold has been predicted to interact with DNA(29), making it an ideal candidate to function in DNA handover between ComEA and the β-lactamase-like domain of ComEC. Therefore, we tested whether the OB fold of *M. glycerini* (hereafter OB_Mg_) can indeed bind to DNA. We performed a fluorescence anisotropy binding experiment where we incubated 12-meric FAM-labelled ssDNA or dsDNA with OB_Mg_ (**Fig. 3a**). This showed that OB_Mg_ binds to dsDNA (dissociation constant (*K*_D_) of 0.33 µM) with approximately 2.4-fold greater affinity than to ssDNA (*K*_D_ of 0.79 µM). Next, we determined the structure of the OB_Mg_ using NMR spectroscopy (**Fig. 3b, Table S2**). The OB_Mg_ structure contains a 6-stranded mixed β-sheet and a 28 residue long disordered loop between β4 and β5. OB family proteins frequently contain small ⍺-helices between β3 and β4 or β4 and β5. However, there are no such helices in our structure, which for the β4-β5 loop could be due to the absence of contacts from the competence domain and/or the β-lactamase-like domain. Although OB fold sequences can be very divergent, both between ComEC sequences as well as between OB folds contained in other proteins, their overall fold is similar (**Fig. S9**). Some OB fold containing proteins, such as the telomere end binding protein, are known to form functional dimers(72), OB_Mg_ behaves as a monomer in solution (**Fig. S10**). Furthermore, the structure of OB_Mg_ does not contain any disulphide bonds, as previously described for the *B. subtilis* OB fold (formerly called the N-loop)(28).

**Fig. 3:**
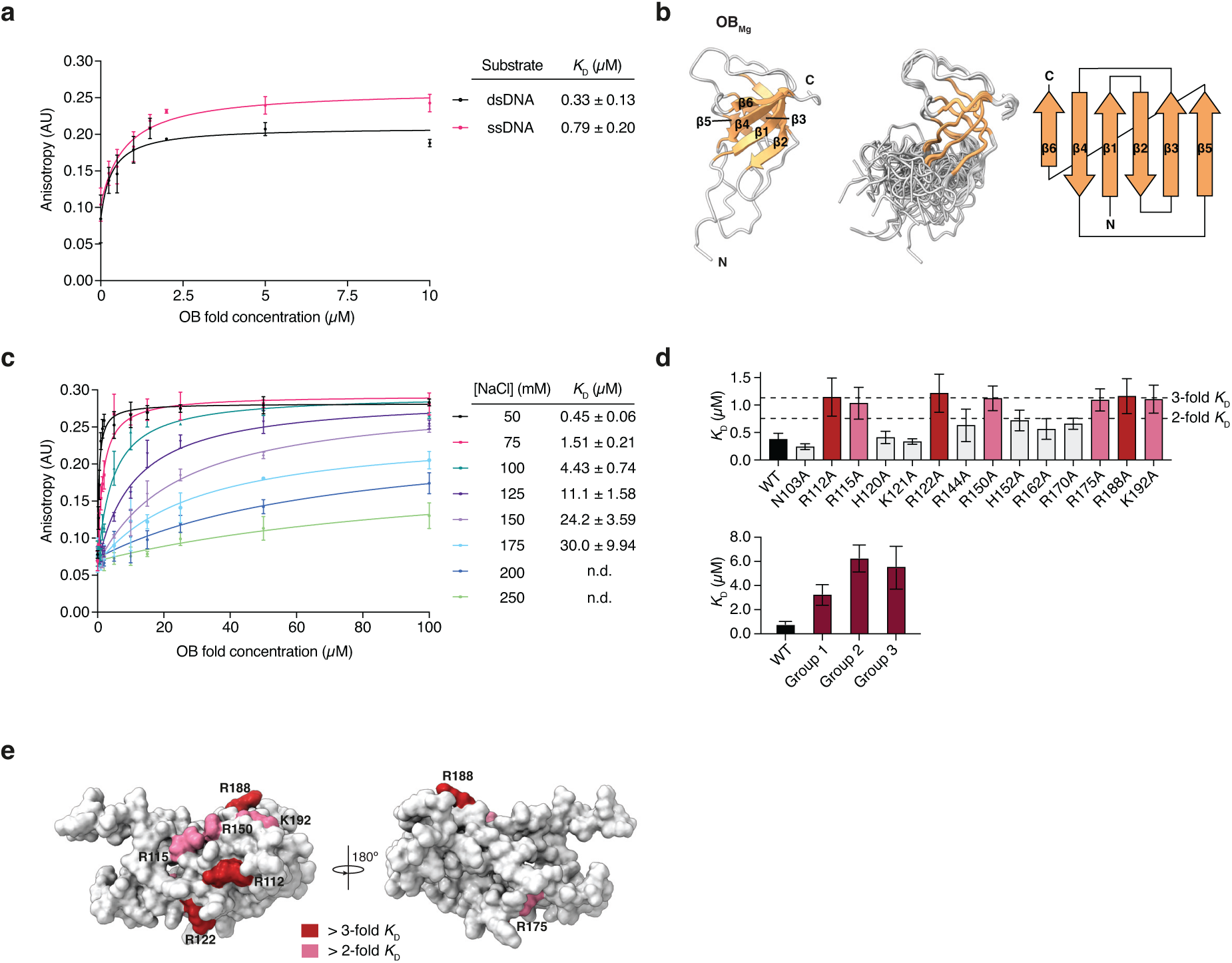
The interaction of OB_Mg_ and DNA occurs through positively charged surface residues and is dependent on ionic strength. **a** Fluorescence anisotropy binding experiments testing the interaction between 12 bp 5′ FAM-labelled ssDNA or dsDNA and OB_Mg_, in a buffer containing 50 mM NaCl. **b** The solution structure of OB_Mg_ shown in ribbon representation (left), a superposition of all states (middle) and the corresponding topology diagram (right). The 6-stranded mixed β-sheet (orange), as well as N- and C-termini are indicated on the figure. **c** Fluorescence anisotropy binding experiments to assess interaction between 12 bp 5′ FAM-labelled dsDNA and wild-type OB_Mg_ in buffers containing 50-250 mM NaCl. All fluorescence anisotropy experiments were performed in triplicate and error bars represent the standard error of the mean. **d** Dissociation constants (*K*_D_) derived from fluorescence anisotropy experiments performed with OB_Mg_ mutants at 50 mM NaCl. Single and triple mutants are plotted separately. For single mutants, thresholds indicating a 2- or 3-fold increase in the mutant *K*_D_ relative to wild-type is marked on the graph by dashed lines. Triple mutants are indicated on the figure as group 1 (R112A/R115A/R122A), group 2 (R150A/ R188A/K192A) and group 3 (R115A/R150A/R188A). WT, wild-type. **e** The fold change in *K*_D_ from (**d**) was mapped onto the surface of OB_Mg_. Residues that increased the *K*_D_ by more than 2- or 3-fold when substituted to alanine are coloured in light pink and red respectively.

We attempted to perform chemical shift perturbation experiments to map the DNA binding interface as well as to further structurally characterise the DNA-bound state by NMR. However, at the high protein concentrations required for these DNA titration or triple resonance experiments, the complex of OB_Mg_-DNA was prone to precipitation. Therefore, we again turned to fluorescence anisotropy binding experiments in order to learn more about the DNA binding mode of OB_Mg_. First, we tested whether the affinity of the interaction between wild-type OB_Mg_ and dsDNA is affected by the ionic strength of the buffer (**Fig. 3c**). Indeed, the *K*_D_ decreased from 30 μM to 0.45 μM, when the NaCl concentration in the buffer was reduced from 175 mM to 50 mM, respectively, suggesting that this interaction is electrostatically driven. Next, we substituted several candidate DNA binding residues to alanine, focusing on positively charged residues, on the surface of OB_Mg_ and repeated the fluorescence anisotropy measurements (**Fig. 3d, Fig. S6**). While some of these single residue substitutions had almost no effect on the affinity of the interaction (e.g. N103A, H120A, K121A), all other single amino acid substitutions increased the *K*_D_ between ∼2-3 fold. We also created triple mutants, where we substituted three nearby residues with alanine, (group 1: R112A/R115A/R122A; group 2: R150A/R188A/K192A: group 3: R115A/R150A/R188A). These triple mutations had a greater effect on DNA binding, resulting in ∼4-19 fold increase in the measured *K*_D_ (given the wild-type *K*_D_ range of ∼0.33 to 0.8 μM). We mapped all residues that, on their own, showed greater than a 2 fold increase in the *K*_D_ onto our NMR structure of OB_Mg_ (**Fig. 3e**). This revealed a binding surface on one side of the molecule that consists of R112, R115, R122, R150, R188 and K192, although the mores structurally isolated R175 also seems to contribute to DNA binding. In the context of the structural prediction of full-length ComEC, this surface is located such that DNA binding to the OB fold and subsequent degradation by the β-lactamase-like domain would be conceivable (**Fig. S11**). Taken together our data show that OB_Mg_ tightly interacts with DNA and that this interaction is electrostatically driven, occurring primarily through several positively charged residues clustered together on the surface of the molecule.

### The OB fold and β-lactamase-like domain of ComEC work in concert for efficient nuclease activity

During natural transformation, competent cells are able to take up vast stretches of DNA(73), Indicating that the uptake machinery must be highly efficient and processive. Yet, throughout this study, we repeatedly noted the relatively poor apparent initial rate of DNA degradation of BLACT_Mg_ (apparent initial rate for ssDNA degradation of ∼0.26 min^−1^) and its seemingly absent affinity for its DNA substrate (*K*_D_ not determinable) (**Table 1, Fig. S12**). As mentioned previously, a close structural homologue of BLACT_Mg_ is RNAse J, which contains an additional β-CASP domain involved in substrate binding. In the structural prediction of ComEC, the OB fold is positioned next to the β-lactamase-like domain (**Fig. S8**). We hypothesised that in a manner analogous to the substrate-binding β-CASP domain of RNAse J, the presence of the OB fold may affect the rate of DNA degradation by the β-lactamase-like domain. As we are currently unable to produce full-length ComEC, we compared the nuclease activity of BLACT_Mg_ on its own and in the presence of OB_Mg_ added in solution or tethered to the BLACT_Mg_ domain as a fusion construct (**Table 1**, **Fig. 4a, b, Fig. S6**). We designed the fusion construct in a way that would mimic the overall ComEC architecture by using maltose-binding protein (MBP) as a scaffold, which is of similar size to the competence domain and would result in similar relative domain positioning according to its predicted structure (**Fig. S13**). These data showed that adding the OB fold in solution did not change the apparent initial rate of DNA degradation significantly (0.26 vs 0.29 min^−1^), whereas in the context of the OB-MBP-BLACT fusion construct the activity was clearly increased (∼7 fold increase in apparent initial rate to 1.85 min^−1^). OB_Mg_ on its own did not display any nuclease activity (**Fig. 4b**). Given that DNA binding by the OB fold is heavily influenced by the ionic strength of the solution (**Fig. 3c**), we tested its effect on the nuclease activity of BLACT_Mg_ and OB-MBP-BLACT (**Table 1**, **Fig. 4c**). Not surprisingly, the ionic strength had a profound effect on the rate of DNA degradation by the OB-MBP-BLACT fusion construct, with the apparent initial rate increasing ∼32 fold from 0.06 min^−1^ to 1.92 min^−1^ when the NaCl concentration was decreased from 500 mM to 50 mM. The effect on the BLACT_Mg_ alone was less pronounced (∼2 fold increase in apparent initial rate), yet still clearly measurable. These data are in agreement with the OB fold serving as the main DNA binding platform for the β-lactamase-like domain. Next, we wondered whether the OB_Mg_ allows for proper substrate positioning with respect to the BLACT_Mg_ active site, or if the simple increase in the local concentration of DNA near the BLACT_Mg_ active site is sufficient to explain the increase in activity. To answer this question we replaced the OB_Mg_ moiety within the fusion construct with another DNA binding protein entirely and performed nuclease activity assays (**Fig. 4d, Fig. S6**). For this purpose we chose the DNA binding domain of ComEA from *M. glycerini* and Sac7d (an OB fold family protein) from *Sulfolobus acidocaldarius*. Like OB_Mg_, these domains are small, soluble and thermostable DNA binding domains that bind to DNA with similar affinities(74, 75). This experiment showed that it does not matter whether the BLACT_Mg_ is fused to OB_Mg_, ComEA_Mg_ or Sac7d, the presence of a DNA binding domain on the same polypeptide chain as the β-lactamase-like domain leads to a similar increase in nuclease activity. This in turn suggests that an increase in local concentration of the substrate is the main mechanism underlying this observed boost in nuclease activity.

**Fig. 4:**
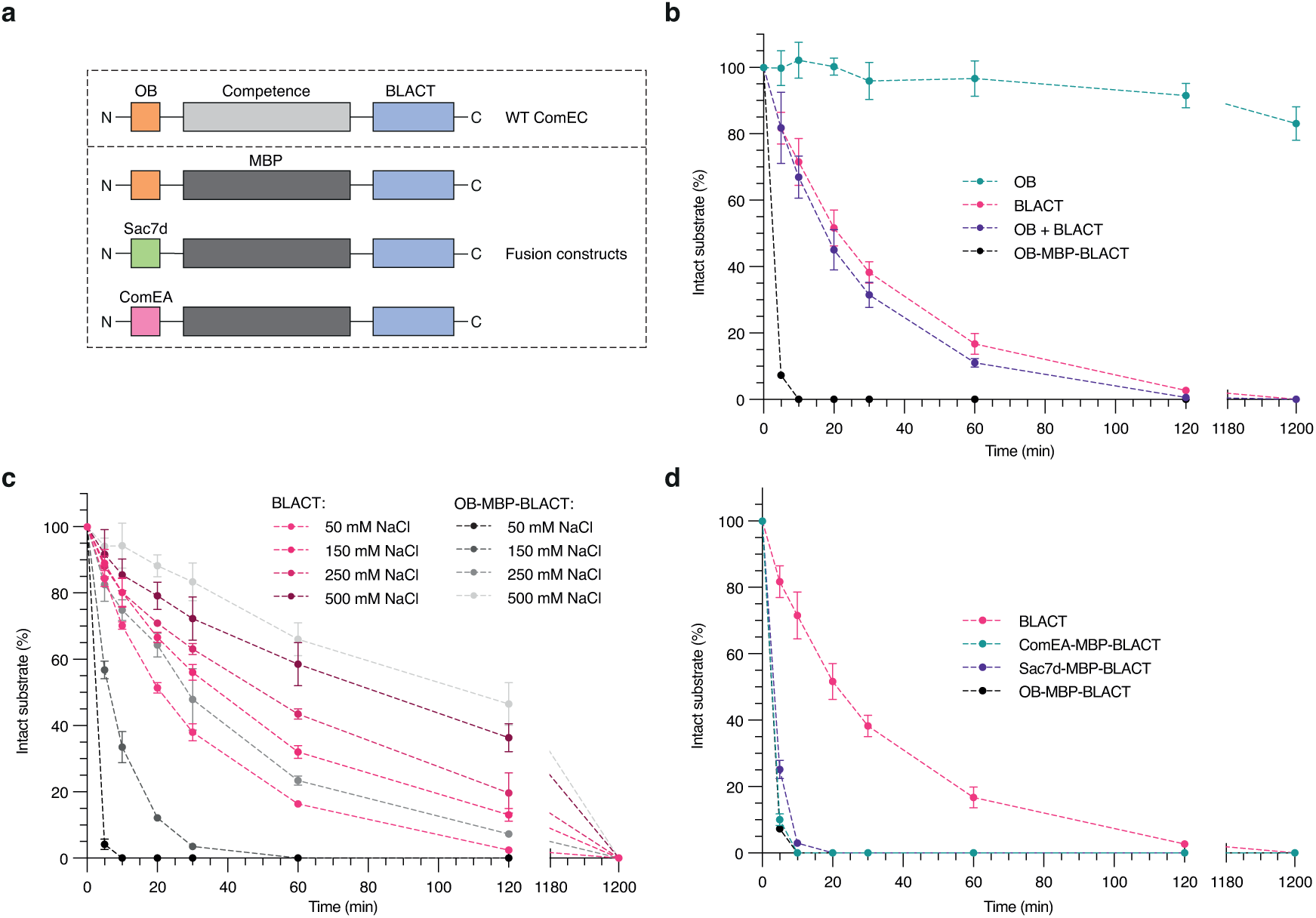
The OB fold serves as the DNA binding domain for the β-lactamase-like domain. **a** Schematic showing the domain organisation of wild-type ComEC and various fusion constructs, encoding either the OB fold or other DNA binding domains (Sac7d from *S. acidocaldarius* and ComEA from *M. glycerini*) on the same polypeptide as the β-lactamase-like domain linked via an MBP scaffold. **b-d** Time-course nuclease activity assay comparing OB, BLACT, OB + BLACT, and OB-MBP-BLACT (**b**), BLACT and OB-MBP-BLACT at four different NaCl concentrations (**c**), and BLACT, OB-MBP-BLACT, Sac7d-MBP-BLACT and ComEA-MBP-BLACT (**d**). Enzyme and substrate concentrations, length of the DNA substrate, and the mode of FAM-labelling of the substrate, were identical to the experiments in Figure 1.

**Table 1:**
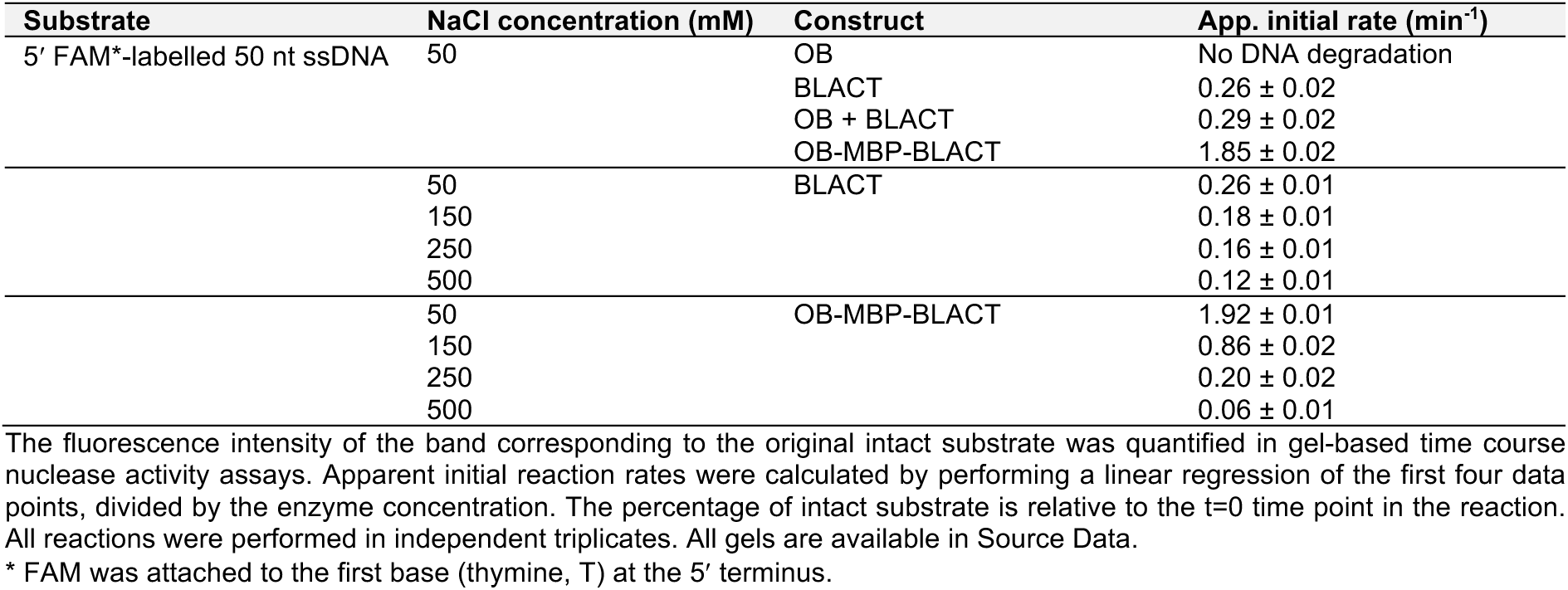
Apparent initial rates of BLACT and OB-MBP-BLACT fusion constructs.

### Key residues in ComEC separate the DNA duplex and guide ssDNA into the β-lactamase-like domain active site or towards the competence channel

As alluded to previously, it is not known whether the β-lactamase-like domain of ComEC encounters ssDNA that has been separated prior to the nuclease step, or dsDNA from which the enzyme is capable of selectively degrading a single strand. It is known however, that most transformation events occur with linear dsDNA as the transforming DNA and our results also show that the OB fold displays slightly higher affinity for dsDNA. We hypothesised that the more likely scenario is that there are structural elements within ComEC that locally destabilise the double helix of dsDNA allowing the β-lactamase-like domain to cleave a single strand, while the undegraded strand is guided towards the channel domain.

To identify putative structural elements and residues that play a role in destabilising the hydrogen bonding of the double helix and guiding of the separated strands, we carefully inspected our BLACT_Mg_ crystal structure and a structural prediction of full-length ComEC in complex with DNA. We also considered that such functionally important residues should be highly conserved. We identified two highly conserved loops, spanning residues 517-527 (loop 1) and 682-694 (loop 2) in *L. pneumophila* ComEC, which could serve as ‘pin elements’ that would destabilise the double helix, akin to pins or wedges of helicase domains(76). Loop 1 contains a tyrosine (Y522) followed by a conserved phenylalanine (F523), while loop 2 contains a conserved arginine (R688) at its tip, followed by two conserved phenylalanines (F689 and F691). These aromatic residues could play a role in denaturing the DNA duplex through ring stacking interactions with the DNA bases and thus also guide the strand destined for degradation (5’ end) towards the active site of the nuclease domain. Moreover, we noticed that there are several other well-positioned aromatic residues, as well as some positively charged residues, that create a pathway between the OB fold and β-lactamase-like domain for the undegraded strand (3’ end) tracking towards the entrance of the channel domain (Y108, Y140, Y154, W212, F331). **Fig. 5a** schematically illustrates the location and putative role of our chosen candidate structural elements and residues. To test whether these residues are important for transformation *in vivo*, we mutated these ComEC residues and performed transformation assays in our *L. pneumophila* Lp02 system (**Fig. 5b**). These results showed that a double mutant in loop 2 (F689A/F691A) completely prevents transformation *in vivo*, whereas a similar double mutant in loop 1 (Y522A/F523A) did not result in decreased transformation. Single mutants on their own did not produce transformation phenotypes, which is consistent with observations of similar pin element single mutants in other systems (e.g. UvrD and PcrA helicases) also not being sufficient to produce a phenotype(77, 78). Mutation of channel lining residues within the OB fold or the competence domain either did not affect transformation, or reduced transformation efficiencies modestly. It is not surprising that most single mutations of channel lining residues do not produce a pronounced phenotype, given the total number of residues that contribute to this aromatically lined DNA tunnel. In summary, we believe that the more conserved loop 2 (hereafter pin loop) is a structural element present within ComEC that locally destabilises the DNA duplex after DNA binding by the OB fold and prior to DNA degradation by the β-lactamase-like domain. Furthermore, a staircase of aromatic and charged residues subsequently guides the ssDNA towards and through the channel domain.

**Fig. 5:**
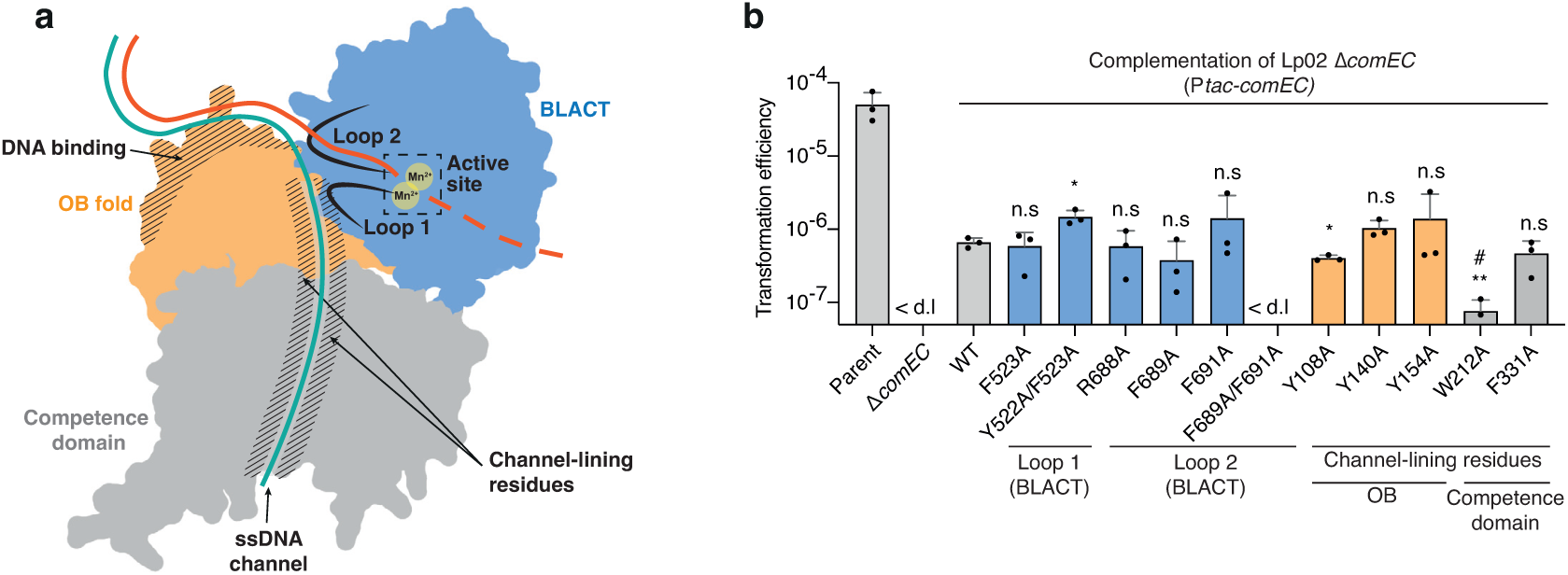
Key residues support DNA degradation and translocation by ComEC. **a** Schematic showing candidate pin and DNA pathway lining residues chosen for mutagenesis. **b** Transformation efficiencies of parental Lp02, Lp02 Δ*comEC*, and Lp02 Δ*comEC* complemented by ectopic expression of wild-type and mutant versions of ComEC containing substitutions of residues in the regions indicated in (**a**). Mean transformation efficiencies of three independent biological replicates are shown with error bars representing the standard deviation (SD). < d.l., below detection limit (d.l.) (average d.l. = 5.09 x 10^−8^). Statistical significance was determined on log-transformed data using an unpaired two-sided t-test with Welch’s correction(116). The Lp02 strain complemented with wild-type ComEC was compared to those strains complemented with ComEC mutants. WT, wild-type; #, below d.l. in at least one replicate; n.s., not statistically significant, *p* > 0.05; *, *p* < 0.05 (*p*_Y522A/F523A_ = 0.01, *p*_Y108A_= 0.02); **, *p* < 0.01 (*p*_F689A/F691A_ = 0.001, *p*_W212A_= 0.006).

### Working model for ComEC

Our *in vitro* characterisation of the OB_Mg_ and BLACT_Mg_, combined with key observations made *in vivo* allow us to propose a working model for DNA binding, degradation and translocation by ComEC (**Fig. 6**). In our model, the OB fold first binds to dsDNA with high affinity. We showed that the residues involved in DNA binding are solvent-exposed and positioned, in the context of the structural prediction of full-length ComEC, in a manner that would seamlessly facilitate DNA capture and subsequent steps (**Fig. S11**). OB fold binding to DNA is necessary as the β-lactamase-like domain on its own does not appear to bind efficiently. Next, structural pin elements locally destabilises the hydrogen bonding between bases of the dsDNA allowing strand separation to occur. Base stacking interactions with the first base pair of the DNA duplex can occur via the conserved aromatic residues present on this loop. The pin loop is located ideally between the ssDNA channel and the active site of the β-lactamase-like domain. Subsequently, the β-lactamase-like domain begins to selectively cleave the strand leading with its 5’ end (**Fig. 2**). The inherent directionality of DNA degradation by the β-lactamase-like domain will thus establish the polarity of DNA translocation through the competence domain. This is in line with previous work that showed the strand leading with the 3’ end is transported into the cytoplasm(79). There is no ATP consumed by ComEC for the translocation of DNA. Given that phosphodiester bond cleavage is energetically favourable and assuming that the β-lactamase-like domain will continue to cleave the 5’ strand, the growing single stranded portion of the 3’ strand is guided by the channel lining residues and threaded into the competence domain. The aromatic and charged residues that line the ssDNA channel likely guide the undegraded DNA strand via ring stacking and electrostatic interactions. Once the DNA emerges on the cytoplasmic side of the membrane, other players like ComFA (and potentially PriA in Gram-negative bacteria), may engage the emerging ssDNA and translocate along it in an ATP-dependent manner, thereby exerting a pulling force. A key feature of our model is the relative domain organisation of ComEC. This defined topology ensures that only a single strand is selectively degraded, as nucleolytic attack of the other strand would require a 180 degree rotation of the β-lactamase-like domain with respect to the DNA, which is presumably held in place through interaction with the OB fold. This topological restraint, coupled with the threading and pulling of DNA through ComEC, theoretically also allows DNA degradation to proceed much more processively *in vivo*, explaining the observed rapid rates of DNA translocation of 80-100 nt/s in *B. subtilis* and *S. pneumoniae*(80, 81).

**Fig. 6:**
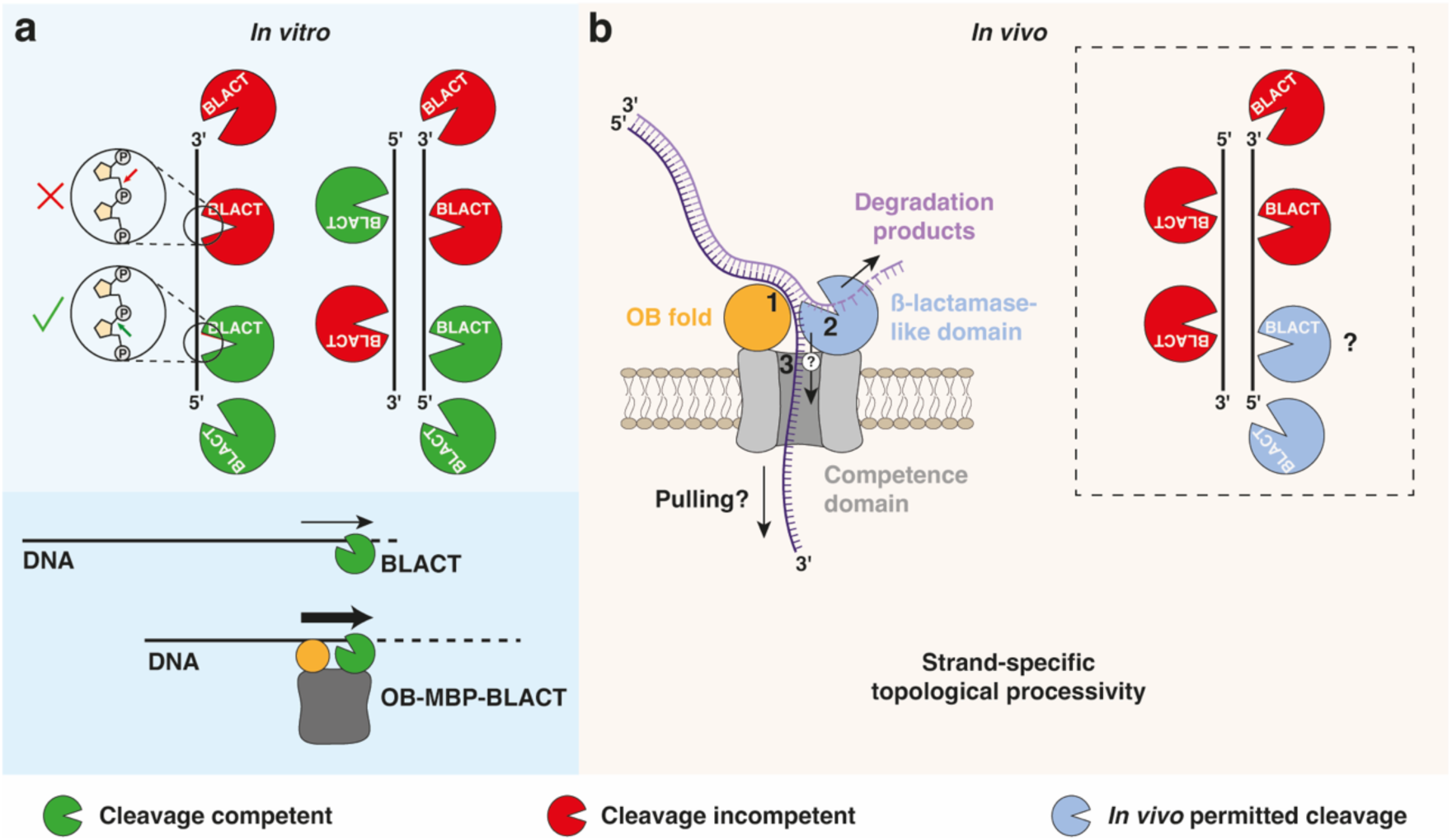
Working model of DNA binding, degradation and translocation by ComEC. **a** Top, possibilities of phosphodiester bond hydrolysis by BLACT_Mg_ *in vitro* shown for both ssDNA and dsDNA. Green shapes represent possible cleavage modes and configurations of BLACT_Mg_, whereas red shapes depict those that cannot occur. All possible cleavage events, including endonucleolytic events, result in a 5′ phosphate and 3′ hydroxyl group as shown in circles. Bottom, the OB-MBP-BLACT fusion construct is able to cleave DNA more efficiently than BLACT alone. **b** Schematic showing key steps carried out by ComEC. 1: tDNA is bound tightly by the OB fold. 2: pin residues on loop 2 locally destabilise the hydrogen bonding between base-paired DNA strands. The 5′ strand is guided towards the nuclease active site by F689 and F691 and 5′-to-3′ exonucleolytic or endonucleolytic cleavage can occur. 3: residues that line the DNA pathway on the OB fold (Y108, Y140 and Y154) and the β-lactamase-like domain (Y522 and F523) guide the DNA towards and through the competence domain. The initial threading of the DNA towards and into the channel domain is likely driven by the activity of the β-lactamase-like domain (depicted by an arrow with a question mark). Once the 3′ strand emerges on the cytoplasmic side of the membrane, other proteins (not shown) can engage with DNA to exert a possible pulling force. The dashed box shows that possible cleavage events by the β-lactamase-like domain, depicted by blue shapes, are reduced *in vivo* due to the topological restraint of ComEC domains and the tDNA.

## Discussion

Natural transformation has far-reaching consequences for bacterial evolution and the emergence of pathogenic strains. Central and essential to the process of DNA transport from the environment into the cell interior is the protein ComEC. Over the years, this protein has proved to be challenging to produce and detect, and as a consequence our mechanistic understanding of this critical step is severely lacking. Here, we characterised the two non-membrane domains of ComEC, the OB fold and the β-lactamase-like domain, to learn more about how ComEC binds to DNA, degrades the non-translocating strand and ultimately translocates the remaining strand. Based on our investigations, we propose a model of how ComEC transports the transforming strand across the cytoplasmic membrane of competent bacteria.

We determined the crystal structure of the β-lactamase-like domain from *M. glycerini* (BLACT_Mg_) (**Fig. 1**). Like its close relative from *M. thermoacetica*(53), this enzyme degrades DNA substrates in a manganese-dependent manner (**Fig. 1c, Fig. S2**). Our structure, in combination with structures of related MBL family proteins, allowed us to propose the catalytic mechanism of this nuclease (**Fig. 1d**). Mutation of key residues that, either prevent Mn^2+^ coordination, or interfere with substrate binding, lead to a reduction or loss of nuclease activity *in vitro* (**Fig. S5**). In our *in vivo* transformation assay, all mutations that reduced nuclease activity *in vitro* completely abrogated transformation, except for S728 which is not involved in metal ion coordination (**Fig. S5**). Such an all-or-nothing phenotype highlights the importance of efficient DNA degradation by the β-lactamase-like domain for successful DNA translocation across ComEC. It remains to be seen, whether the β-lactamase-like domains from other competent species are dependent on Mn^2+^ coordination, or whether some of these enzymes may indeed bind to other metal cofactors such as Zn^2+^, as was predicted in earlier studies(29).

Nucleases often possess both endo- and exonuclease activity(82). A close structural homologue of BLACT_Mg_ is RNAse J, which can also cleave its RNA substrate in both modes, and indeed switch its propensity towards one mode depending on the nature of the 5’ modification of RNA or the enzyme’s dimerisation status(61, 62). We investigated whether BLACT_Mg_ shows an intrinsic preference towards cleaving single- or double-stranded DNA substrates, whether or not the presence of a 5’ phosphate or hydroxyl group is preferred and the directionality of its exonuclease activity (**Fig. 2**). These results showed a clear preference for ssDNA and revealed that BLACT_Mg_ can also hydrolyse in both an endo- and exonucleolytic fashion, the latter occurring exclusively in the 5’-to-3’ direction. This would suggest that when the β-lactamase-like domain encounters DNA *in vivo*, that the strand leading with its 5’ end is degraded, while the 3’ strand is free to translocate. Therefore, the observed 3’-to-5’ directionality of DNA translocation through ComEC(79) can be explained mechanistically by the enzyme’s inherent directionality of DNA degradation.

Whether or not both endo- and exonuclease activities are required during DNA translocation *in vivo* remains to be further investigated. However, we believe it is conceivable that the β-lactamase-like domain could operate in a mixed exo- and endonucleolytic manner, and that such endonucleolytic cleavage events may occur sporadically as the enzyme skips one or several bonds once the tDNA has become engaged inside ComEC, thereby releasing units that are longer than a single nucleotide. Structurally, there is sufficient solvent space around the active site to permit endo-cleavage events that would result in the release of such longer units. Indeed, there is evidence for this from earlier work that showed that di- and tri-nucleotides are released during this process in *B. subtilis* and *S. pneumoniae*(83). Alternatively, the endonuclease activity might serve to create new DNA ends that can be engaged by ComEC, however, this seems less likely as it would require an attack on the opposite DNA strand that would be difficult to envision given that the β-lactamase-like domain is not free to rotate in solution.

Curiously, BLACT_Mg_ on its own does not display measurable DNA binding activity (**Fig. S12**), and our data show that the OB fold provides this critical function (**Fig. 3**). OB_Mg_ binds to DNA tightly (*K*_D_ = 0.33 µM for dsDNA) using several positively charged residues as demonstrated by the drastic effect of ionic strength on binding. Yet substitutions of residues that contribute to binding (single and triple alanine substitutions) do not reduce affinity greatly. It appears as though the OB fold of ComEC has evolved as a robust DNA binding module that cannot easily be perturbed by single amino acid substitutions.

In the absence of an auxiliary DNA binding domain such as a β-CASP domain, the OB fold thus fulfils this function (**Fig. 4**). This was demonstrated by the ∼7 fold increase in the apparent initial rate of DNA degradation when OB_Mg_ was fused to BLACT_Mg_ on the same polypeptide chain and the dependence of DNA degradation on ionic strength, mirroring the OB_Mg_-DNA binding experiments. It does not appear that OB_Mg_ plays a role in the correct positioning of the substrate with respect to the BLACT_Mg_ active site in our fusion construct *in vitro*, since the increased DNA degradation activity can also be achieved by other DNA binding domains. However, at present we cannot determine if the precise domain orientation within the native ComEC protein *in vivo*, could potentiate activity further still. Structural alignment of the MBL domains within RNAse J and the predicted structure of full-length ComEC reveal the different relative positions of the OB fold and the β-CASP domain with respect to the nuclease domain (**Fig. S14**), suggesting that the route taken by the RNA/DNA substrate to the nuclease domain active site differs between these two proteins. In RNAse J from *S. coelicolor* (PDB ID: 5A0T)(61) and *Deinococcus radiodurans* (PDB ID: 4XWW)(62) the 5’ phosphate of the RNA substrate is coordinated by a conserved serine (S375 and S379, respectively). The S728A variant of BLACT_Mg_ (S662 in *L. pneumophila*) showed reduced nuclease activity (**Fig. S5**), presumably because of its role in substrate coordination. However, in the absence of a substrate bound complex structure and due to the different positioning of the OB fold relative to the β-lactamase-like domain within ComEC compared to the domain arrangement of RNAse J, it is difficult to precisely pinpoint further substrate coordinating residues. Yet it is precisely this alternative placement of the OB fold within ComEC that likely creates the DNA tunnel that will ultimately guide the non-degraded ssDNA towards the competence domain.

To more precisely understand how DNA might be bound and subsequently encountered by the active site of the β-lactamase-like domain, we performed mutagenesis and transformation experiments guided by structural predictions and our own data (**Fig. 5**). We identified a structural element, the pin loop, that separates dsDNA prior to degradation. We showed that substitution of two conserved phenylalanines (F689A and F691) on this loop completely abrogates transformation. This ability of ComEC to locally melt and separate incoming duplex DNA is a critical aspect of our working model and is reminiscent of the mechanisms employed by helicases(76, 77), as well as other DNA processing proteins such as T7 RNA polymerase(84), which use similar structural elements. Initially this process in ComEC is not powered by ATP hydrolysis, but is likely driven by the activity of the β-lactamase-like domain. Because one of the two DNA strands is immediately degraded, no torsional backpressure or overwinding problems can occur. As cleavage progresses, the ssDNA portion of the strand destined for translocation ‘grows’, which then takes the path of least resistance and is threaded into the competence domain. To this end, the relative domain organisation of OB fold, β-lactamase-like domain and competence domain ensure that a clear pathway is established and the DNA is prevented from taking an alternative route. The channel lining aromatic residues help to guide the ssDNA through this pathway. Indeed, the putative DNA channel within ComEC is lined with several well-positioned and highly conserved aromatic, as well as some charged, residues (**Fig. 5, 6**), which is a hallmark of proteins that bind to, stabilise or translocated DNA in some manner. For example, such a helical gateway is observed in the RecJ nuclease, which is important for processively degradation of ssDNA by this enzyme(85). Interestingly, this protein also contains an OB fold which is important for DNA binding, and its relative position with respect to the nuclease seems equally critical. In the case of natural transformation, once ssDNA emerges on the cytoplasmic side of the membrane and is engaged by an ATP-dependent DNA translocase (ComFA/PriA), unwinding, degradation and translocation may be further accelerated.

Many proteins involved in processive reactions of RNA or DNA metabolism achieve processivity via the topological linkage model, where ring-shaped, oligomeric proteins or protein complexes encircle their linear nucleic acid substrate (examples(86–90)). Among these examples are helicases, the sliding clamp β (a processivity factor of DNA polymerase III) and lambda exonuclease. The latter is a homotrimeric enzyme with a central DNA channel that processively cleaves one strand of a dsDNA substrate by virtue of its funnel-shaped channel that is wide enough to encircle dsDNA at one end, but can only fit ssDNA at the other(90). In our model of ComEC, the relative domain organisation imparts a topological restraint to the system, which we believe is crucial for the β-lactamase-like domain to cleave DNA processively *in vivo* (**Fig. 6**). We term this strand-specific topological processivity. Since the β-lactamase-like domain cannot diffuse and rotate 180°, endonucleolytic attack of the translocating strand leading with its 3’ end cannot occur, thus sparing it from degradation. This allows the nuclease domain to engage and processively cleave the 5’ strand. Furthermore, the uncleaved strand is subsequently stabilised and guided towards the channel domain of ComEC. According to structural predictions, this channel would not be wide enough to accommodate dsDNA. Further work is required to precisely understand the energetics of DNA duplex separation, DNA degradation and ssDNA translocation.

In summary, our work allows us to propose a working model for how ComEC binds, degrades and translocates DNA through the membrane during transformation. This is an important step towards developing a complete mechanistic model of this essential protein, which has until now remained poorly characterised.

## Methods

### Bacterial strains and growth conditions

*L. pneumophila* Lp02 strains (derived from *L. pneumophila* Philadelphia 1) were cultured in ACES [N-(2-acetamido)-2-aminoethanesulfonic acid] buffered yeast extract (AYE) liquid medium. For growth on solid medium, ACES-buffered charcoal yeast extract (CYE) supplemented with 100 μg/mL streptomycin and 100 μg/mL thymidine (CYE ST) was utilised. All media additionally included 0.4 g L-cysteine, 0.135 g Fe(NO_3_)_3_ per litre of culture. For selection, 20 μg/mL kanamycin or 5 μg/mL chloramphenicol were added when appropriate. **Table S3** provides a list of all bacterial strains used in this study.

### Plasmids

All constructs for recombinant protein expression were generated with the pOPINS vector(91). The vector contains an N-terminal His_6_-SUMO tag and inserts were cloned in frame downstream of the T7 promoter. Template DNA of the ComEC and ComEA genes from *M. glycerini* and the Sac7d gene from *S. acidocaldarius* were synthesised (Twist Bioscience) prior to further cloning. Independent constructs of OB_Mg_ (residues 76-199) and BLACT_Mg_ (residues 532-797) and various fusion constructs were created. The fusion constructs encoded either OB_Mg_ (76-199), Sac7d_Sa_ (1-66) or ComEA_Mg_ (147-211) on the same polypeptide as BLACT_Mg_ (532-797), linked via an MBP scaffold (OB_Mg_-MBP-BLACT_Mg_, Sac7d_Sa_-MBP-BLACT_Mg_, ComEA_Mg_-MBP-BLACT_Mg_). GSSGSS linker sequences were introduced between DNA binding domain and MBP, and MBP and BLACT_Mg_. Constructs for *in vivo* transformation assays were generated with pMMB207C by insertion of relevant constructs downstream of the P*tac* promoter(92). In-Fusion cloning and site-directed mutagenesis were carried out using the CloneAmP HiFi PCR premix (Takara) according to the manufacturer’s instructions. All plasmids used in this study are listed in **Table S4**, while primer sequences can be found in **Table S5**.

### Protein production

All proteins were N-terminally His6-SUMO tagged and expressed in BL21 (DE3) *E. coli* cells. Cultures were grown in Luria-Bertani (LB) media at 37°C until an optical density at 600 nm (OD_600_) of 0.6-0.8 was reached, while shaking. Cultures were then induced with 0.5 mM β-D-thiogalactoside (IPTG) and further incubated for 12-18 h at 18°C, while shaking. Cells were lysed in 50 mM HEPES-NaOH pH 7.2, 1 M NaCl, 40 mM imidazole, supplemented with 0.2 mg/mL lysozyme, 10 µg/mL DNAse, and one complete mini EDTA-free protease inhibitor tablet (Roche). Cells were lysed by passing the suspension three times through an EmulsiFlex-C5 homogeniser (Avestin) at 40000 psi. The lysate was clarified by centrifugation in a JLA-16.250 (Beckman Coulter) at 30’000 *g* for 60 min, filtered through a membrane with a pore size of 0.22 µm and applied to a 5 ml HisTrap HP column (Cytiva). Elution was performed either with a linear 40-500 mM imidazole gradient or by a stepwise elution with 500 mM imidazole. Protein containing fractions were pooled and dialysed against 50 mM HEPES-NaOH pH 7.2, 50 mM NaCl, while the His_6_-SUMO tag was cleaved by addition of the catalytic domain of the human SENP1 protease to the dialysate. The OB_Mg_ and BLACT_Mg_ were further purified by ion exchange chromatography using a 5 mL HiTrap Q HP column (Cytiva), collecting the protein of interest in the unbound fraction. In contrast, fusion proteins were purified using a 5 mL HiTrap SP HP column (Cytiva), eluting bound proteins using a linear salt gradient from 50 mM to 1 M NaCl. The final purification step was size exclusion chromatography of the samples using either a HiLoad 16/600 Superdex 75 pg column or a 10/300 GL increase 75 pg column (Cytiva). Protein solutions were concentrated using centrifugal filter devices with a molecular weight cut-off of 10 or 30 kDa (Millipore) and the concentration was determined by measuring the specific absorption at 280 nm, using the molar extinction coefficient of 14900 M^−1^ cm^−1^ for OB_Mg_, 26930 M^−1^ cm^−1^ for BLACT_Mg_, 108180 M^−1^ cm^−1^ for OB-MBP-BLACT, 101760 M^−1^ cm^−1^ for Sac7d-MBP-BLACT and 94770 M^−1^ cm^−1^ for ComEA-MBP-BLACT. All purification steps were performed at room temperature, except overnight tag cleavage, which occurred at 4°C.

### X-ray crystallography

BLACT_Mg_ was crystallised using the sitting drop vapour diffusion method at 20°C at a concentration of 10 mg/mL in 30% (w/v) precipitant mix 1 (PEG 500 MME, PEG 20’000), 0.1 M buffer system 1 (1M MES and 1M imidazole mixed in 56:44 ratio to achieve pH 6.5), 0.09 M NPS mix (0.3 M sodium phosphate dibasic dihydrate, 0.3 M ammonium sulfate, 0.3 M sodium nitrate) (well C1, Morpheus I, Molecular Dimensions). Diffraction data were collected at the Swiss Light Source (SLS) beamline X10SA (PXII) at a wavelength of 0.999989 Å. Data processing was performed within the CCP4i program suite(93, 94). The data were indexed and scaled using iMOSFLM(95) and AIMLESS(96), respectively. There is one molecule in the asymmetric unit and the crystal belongs to the space group C 2 2 2_1_. The structure was determined by molecular replacement in MOLREP(97) using an AlphaFold2-generated search model lacking any active site metal ions. The protein chain and phosphate group were iteratively built in COOT(98) and refined in REFMAC5(99) and PHENIX(100). The refinement strategy included positional refinement, solvent correction and individual B-factor refinement. Final statistics for the BLACT_Mg_ structure can be found in **Table S1**.

### NMR spectroscopy

#### Production of isotope-labelled OB_Mg_

Uniformly ^13^C, ^15^N-labelled OB_Mg_ was produced by growing cells in M9 minimal medium containing 1 g/L ^15^NH_4_Cl and 3 g/L ^13^C_6_-glucose, supplemented with 2 mM MgSO_4_, trace elements, vitamin mix and 50 μg/mL kanamycin for selection. Protein expression and purification were performed as described above.

#### Data acquisition and structure determination

For NMR resonance assignments and structure determination, samples consisting of 1.5 mM uniformly ^13^C, ^15^N-labelled OB_Mg_ in 50 mM HEPES, pH 7.2, 50 mM NaCl and 10% D_2_O were used. Spectra were recorded at 25 °C in 3 mm diameter NMR tubes (Bruker). 3D HNCACB(101, 102) and 3D CBCACONH(103) spectra were recorded on a 700 MHz AVNEO spectrometer equipped with a TCI cryo-probe (Bruker). The spectra consisted of 2048×50×90 complex points in the ^1^H, ^15^N, and ^13^C dimensions with respective spectral widths of 16, 34, and 64 ppm and were recorded with 8 scans per increment resulting in 2 and 1.5 days of measurement time, respectively. Side chain assignments were hampered by the intense signals of the HEPES buffer and therefore an alternative sample with deuterated Tris (d-Tris) (Sigma, 449105) was produced. With d-Tris as a buffer substance however, the protein could not be concentrated to the same level as in HEPES buffer. The concentration was sufficient for a 3D HcC(aliaro)H-TOCSY spectrum(104) recorded on a 600 MHz AVIIIHD spectrometer equipped with a TCI cryo-probe (Bruker). The spectrum consisted of 1536×75×150 complex points in the ^1^H, ^1^H, and ^13^C dimensions with respective spectral widths of 16, 10, and 140 ppm, and was recorded with 2 scans per increment in 3 days using a recycle delay of 2 s. NOESY spectra were recorded for both types of samples: the sample in d-Tris buffer produced a clean NOESY spectrum, however, with limited sensitivity, and the sample in HEPES buffer produced a highly sensitive spectrum where the region between 2.8 and 3.9 ppm could however not be interpreted due to strong T_1_ noise. In detail, time shared 3D [^13^C/^15^N, ^1^H]-HSQC NOESYs (modified from(105)) were recorded on a 900 MHz AVNEO spectrometer equipped with a TCI cryo-probe (Bruker). The spectra consisted of 1536×120×256 complex points in the ^1^H, ^1^H, and ^13^C/^15^N dimensions with respective spectral widths of 16, 11, and 140/80 ppm, and were recorded with 2 scans per increment in 3 days.

Resonance assignments were determined with the program cara (www.cara.nmr.ch). The signal intensities exhibited strong variations and only stretches including amino acid residues 11–80 and 107–124 could be assigned to 96% completeness (Figure S2). Automated peak picking of NOESY spectra was performed with the program ARTINA(106) and peak lists were manually cleaned from artefacts using the ccpnmr 2.5.1 software package(107). Resonance assignments and peak lists from both samples were combined and were used as input for a structure calculation with ARTINA and CYANA (version 3.98.15(108)). 140 angle constraints were automatically generated from Cα chemical shifts, and 2098 unambiguous NOE distance restraints were used to calculate a bundle of 100 conformers, from which the 30 with the lowest CYANA target function were selected for refinement in implicit water in the program amber20(109) and the final 20 with the lowest energy were used to represent the structure. A total of 43 hydrogen bonds was identified in more than six structures, Ramachandran plot statistics were as follows: 92.1 %, 7.7 %, 0.1 % and 0.1 % in favored, allowed, generously allowed and disallowed regions, respectively, as defined by the program Procheck(110). Further structural statistics can be found in Table S2.

### Nuclease activity assays

Various DNA probes were tested in nuclease activity assays (Microsynth, **Table S5**). A fluorescein (FAM)-label was attached to a thymine (T) base either at the 5’ or 3’ terminus of all substrates, and some substrates contained PTO bonds. To generate linear FAM-labelled dsDNA, a single strand of FAM-labelled DNA was annealed with the complementary unlabelled strand. Plasmid DNA, circular ssDNA and dsDNA, was obtained from ThermoFisher, Takara, and Microsynth, respectively. Reactions were performed by mixing either 10 μM linear ssDNA (50 nt), 5 μM linear dsDNA (50 bp), 27 nM circular ssDNA (M13mp18, 7429 nt) or 22 nM circular dsDNA (pBR322, 4361 bp) with 1 μM enzyme in 50 mM HEPES-NaOH pH 7.2, 50 mM NaCl, 5 mM MnCl_2_, in a total volume of 100 μL. Reactions were incubated at 50°C in a TAdvanced thermocycler (Biometra) and timepoints were taken by removing 5 μL of the reaction and quenching it with 5 μL Novex 2X TBE-UREA sample buffer (Invitrogen). Samples were resolved on 12% polyacrylamide gels containing 7 M urea and fluorescence detection was achieved using a ChemiDoc imaging system (Bio-Rad).

Fluorescent band intensities were measured (GelAnalyzer V19.1) and normalised to the t=0 time point (corresponding to either 5 μM dsDNA or 10 μM ssDNA) and the percentage of intact substrate remaining was plotted against time. All measured fluorescence intensity values were within a linear range, as confirmed by a standard curve of known DNA concentrations. The apparent initial reaction rate was determined by linear regression of the first four data points, divided by the enzyme concentration. The plasmid DNA degradation experiments were resolved on a 1% (w/v) agarose gel and visualised using UV illumination (Carestream). Each reaction was performed and analysed at least three times. Experiments for Fig. 1c, Fig. 2a, Fig. 4b, d, and Fig. S5a were performed at the same time, therefore the data for BLACT are the same across these panels. Additionally, data for OB-MBP-BLACT are the same in Fig. 4b, d. **Table S5** provides a list of all oligonucleotide substrates used for biophysical assays

### Fluorescence anisotropy

All DNA binding experiments were performed using either 12 bp FAM-labelled dsDNA, generated by annealing a 12-meric 5′ FAM-labelled strand and a complementary unlabelled strand or 12-meric 5′ FAM-labelled ssDNA. In some experiments, all phosphodiester bonds were replaced by nuclease-resistant PTO bonds (Microsynth, **Table S5**). In contrast to the nuclease activity assays, here, the FAM moiety was attached to the 5′ phosphate of the oligo. A constant DNA concentration of 20 or 50 nM was incubated with increasing concentrations of protein in 50 mM HEPES-NaOH pH 7.2, 50 mM NaCl for 60 minutes at 25°C. To test the effect of ionic strength on binding, some experiments were conducted with buffers containing different NaCl concentrations, as indicated in the relevant figures. Following incubation, 30 μL of each sample was placed in a 96-well half area black flat bottom polystyrene plate with a non-binding surface (Corning). Fluorescence intensities, parallel and perpendicular to the excitation polarisation, were measured in a Synergy2 plate reader (BioTek) (excitation: 495 nm; emission: 520 nm). The anisotropy was calculated using the following equation (Eq. 1),

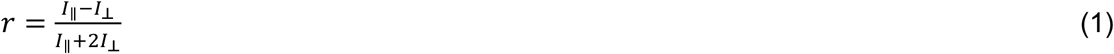

where r is the anisotropy, I|| is the fluorescence intensity in the parallel direction, and I_┴_ is the fluorescence intensity in the perpendicular direction. The calculated anisotropy for each sample was plotted against the protein concentration and the curve was fitted to a model assuming one set of binding sites in order to derive the dissociation constant (*K*_D_) (Eq. 2),

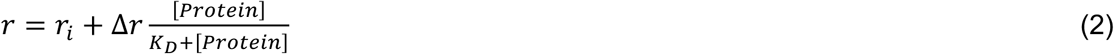

All binding measurements were performed at least three times. As an additional control, some samples were measured twice, 60 min apart, to ensure that the binding equilibrium was fully attained at the time of measurement.

### Transformation assays

Transformation assays were performed as previously described(15). Frozen stocks or freshly transformed Lp02 cells were streaked onto CYE plates supplemented with appropriate antibiotics, and incubated at 37°C for 3-5 days until colonies appeared. Bacteria were resuspended in a 10 mL AYE liquid starter culture and incubated overnight at 37°C while shaking. Fresh 10 mL AYE cultures were inoculated with overnight starter cultures and incubated at 30°C while shaking. At an OD_600_ of 0.3, 1 mL of the culture was transferred to a new tube, mixed with 1 µg of tDNA, and further incubated at 30 °C for 24 h. The tDNA fragment, containing kanamycin resistance cassette, was described previously(15, 111). For complementation experiments, 0.5 mM IPTG was added simultaneously with the tDNA to induce ectopic expression of genes under the control of an IPTG-inducible P*tac* promoter. Serial dilutions of the culture were spread onto selective and non-selective plates and colony forming units (CFUs) were counted after 4 days of incubation at 37°C. The final transformation efficiency was calculated by dividing the number of CFUs on selective plates by the number of CFUs on non-selective plates. Plates containing fewer than 10 CFUs were not counted.

### Bioinformatic analyses

#### Sequence alignments and conservation analysis

A set of 2000 ComEC sequences were retrieved by BlastP (Blast v2.15.0) against the full-length *M. glycerini* ComEC sequence using a 95% query coverage cutoff. All searches were performed with the refseq_select database, which contains only one reference genome for each prokaryotic species to reduce redundancy in the search. The search was conducted with the Gram-positive exclusion filter followed by an identical search with the Gram-negative exclusion filter to ensure even selection of ComEC sequences from Gram-positive and Gram-negative organisms. The list of sequences was further curated manually by removing any redundant sequences. A multiple sequence alignment was performed with ClustalOmega(112). Conservation was mapped onto the predicted structure of full length-ComEC, the BLACT_Mg_ crystal structure, and the OB_Mg_ NMR structure using the ConSurf server(113).

#### 3D protein structure prediction and comparison

The AlphaFold3 server (https://www.alphafoldserver.com) was used to predict structural models of full-length ComEC_Mg_, ComEC_Lp_, and the OB_Mg_-MBP-BLACT_Mg_ fusion construct(114). To identify structural homologues of BLACT_Mg_ and OB_Mg_, their PDB files were submitted to the DALI protein structure comparison server (http://ekhidna2.biocenter.helsinki.fi/dali/) and hits with high Z-scores were chosen for further comparison(66). Structural figures were produced in ChimeraX(115).

## Data availability

The data that support the findings of this study are available from the corresponding author upon reasonable request. Coordinates and structure factors for BLACT_Mg_ have been deposited in the Protein Data Bank (PDB ID: XXXX). NMR spectra and corresponding model coordinates of OB_Mg_ have been deposited in the BioMag Resonance Data Bank (BMRB: XXXXX) and Protein Data Bank (PDB ID: XXXX), respectively. Source data are provided with this paper.

## Acknowledgements

This work was funded by an SNSF PRIMA grant PR00P3_179728 to MKH. We are grateful to R. Glockshuber, E. Weber-Ban for helpful discussions and sharing of instruments. We thank Stefanie Holz for the pMMB207C-*comEC_Lp_* construct. The graphical abstract was partially created in https://BioRender.com.

## Author Contributions

MJMS, SD, SAGB and DW cloned constructs, purified proteins and performed bioinformatic analyses. SAGB, DW, MJMS and SD carried out nuclease activity assays. MJMS and MGB performed fluorescence anisotropy experiments. SD and SAGB carried out transformation assays. MJMS determined the crystal structure of BLACT_Mg_ and DW performed the final rounds of model building and refinement. ADG conducted and analysed all NMR-related experiments with the help of MJMS. MKH designed and supervised the study and wrote the manuscript with help from all authors. All authors contributed to figures.

