## Supplementary information for "Molecular interplay between ComEC domains leads to efficient DNA translocation during natural transformation"

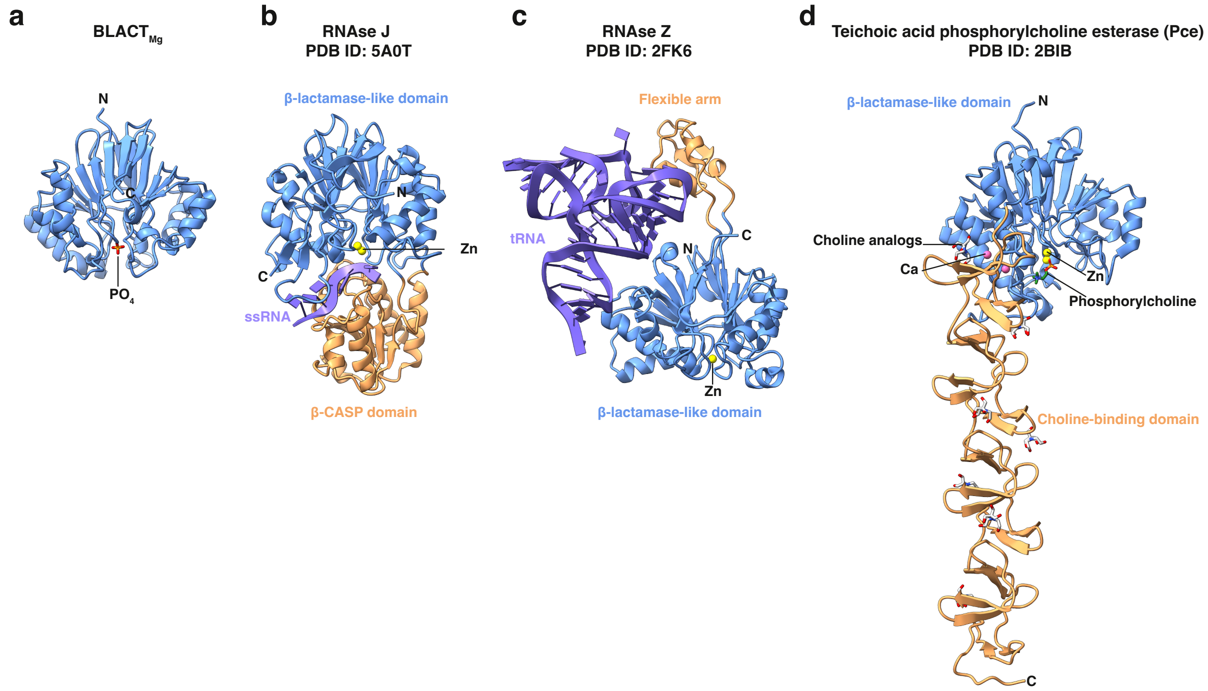

**Fig. S1: Structural comparison of BLACT$_{Mg}$ homologues**

**a-d** Structures of the β-lactamase-like domain from *Moorella glycerini* (BLACT$_{Mg}$) (this study) (**a**), RNAse J from *Streptomyces coelicolor* (PDB ID: 5A0T)[1] (**b**), the RNAse Z/tRNA complex from *Bacillus subtilis* (PDB ID: 2FK6)[2] (**c**), and teichoic acid phosphorylcholine esterase (Pce) from *Streptococcus pneumoniae* (PDB ID: 2BIB)[3] (**d**). Structures are displayed in ribbon representation with the β-lactamase-like domains coloured in blue, substrate binding domains in orange, and RNA molecules in purple. The zinc and calcium ions are shown as yellow and pink spheres, respectively. All ligands are labelled and shown in stick representation.

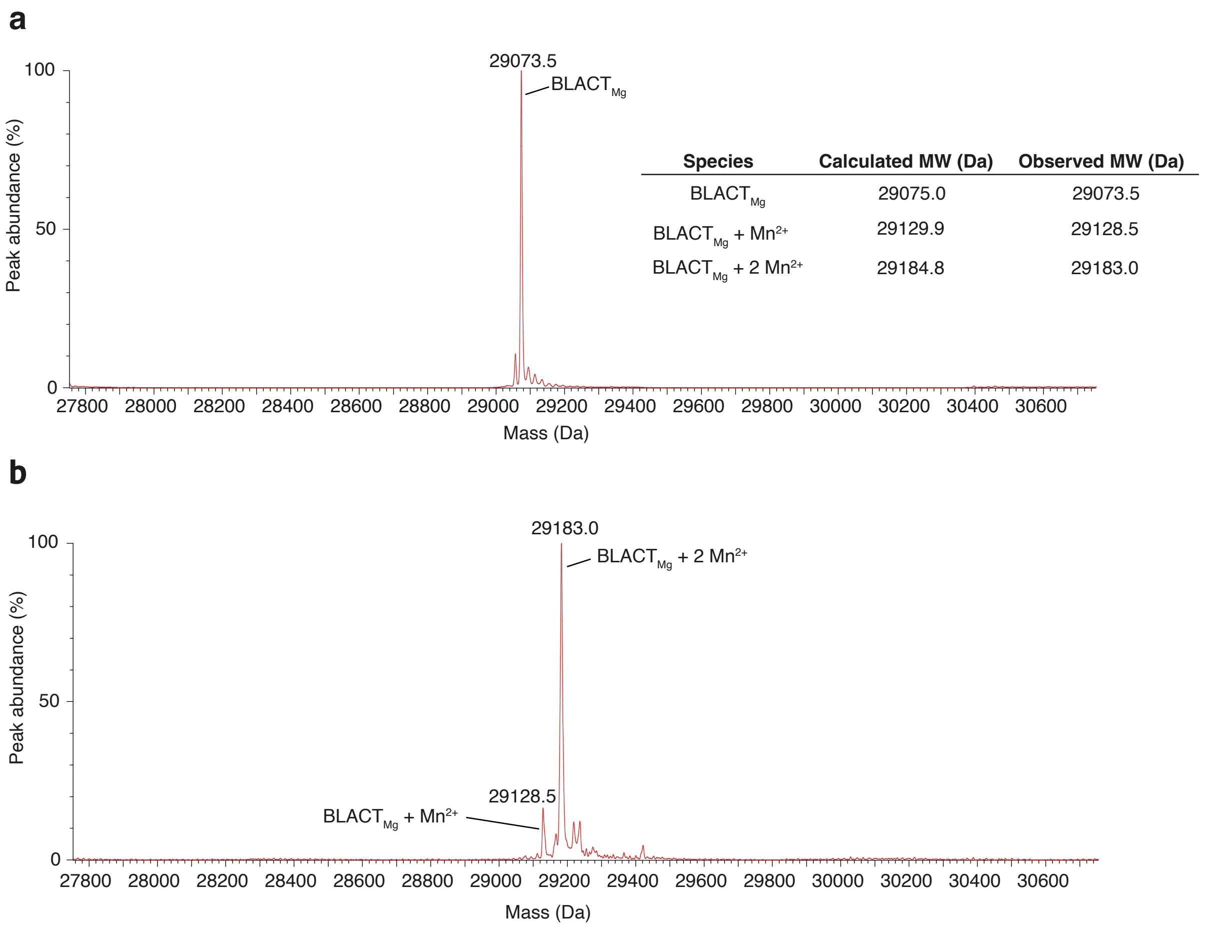

**Fig. S2: The active site of $BLACT_{Mg}$ can coordinate two manganese ions**
**a, b** Intact mass measurements of $BLACT_{Mg}$ without supplemented ions (**a**) and in the presence of 5 mM $MnCl_2$ (**b**). A table of theoretical molecular weights (MW) is shown as an inset. In the absence of supplemented $MnCl_2$, a peak corresponding to the expected mass of $BLACT_{Mg}$ alone is observed. Two peaks are observed in the presence of $MnCl_2$ corresponding to expected masses of $BLACT_{Mg}$ + $Mn^{2+}$ and $BLACT_{Mg}$ + 2 $Mn^{2+}$. The peak corresponding to the fully coordinated state (+ 2 $Mn^{2+}$) is the most abundant peak.

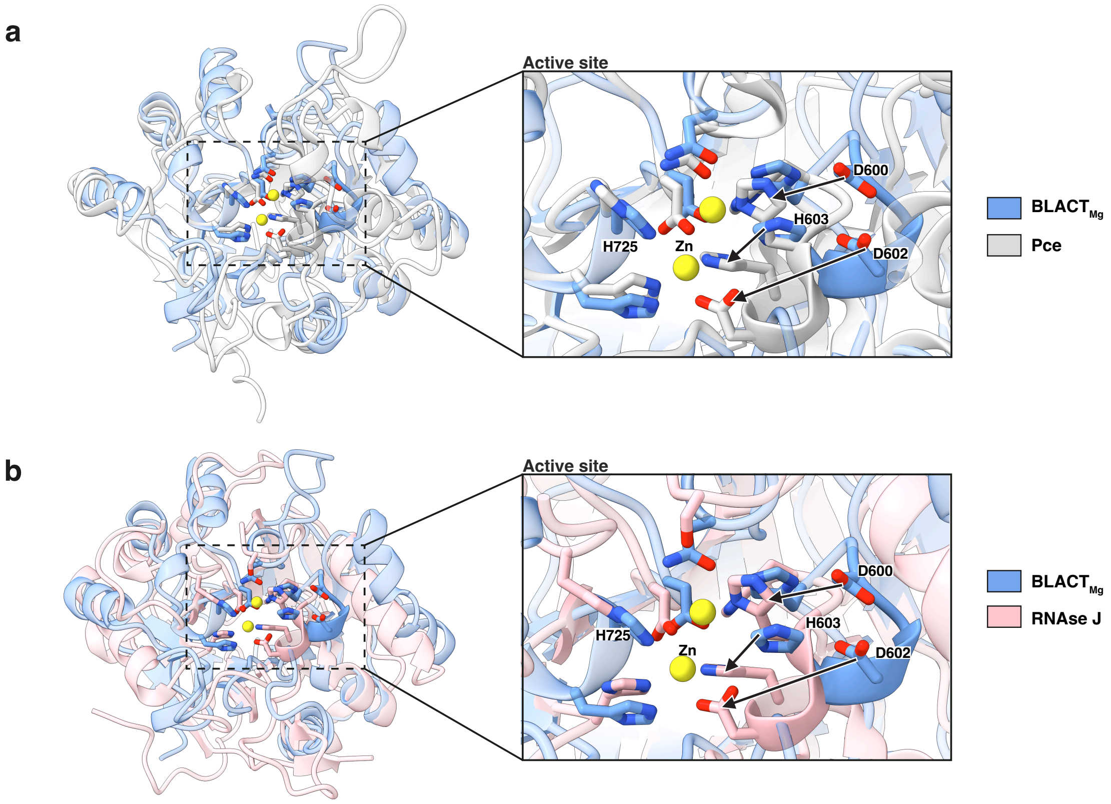

**Fig. S3: Structural differences between the active sites of BLACT$_{Mg}$ and the structural homologues Pce and RNAse J**

**a** Superposition of BLACT$_{Mg}$ (blue, this study) and teichoic acid phosphorylcholine esterase (Pce) from *S. pneumoniae* (grey, PDB ID: 2BIB)[3]. Structures are displayed in ribbon representation, where the side chain of key residues are shown as sticks and Zn ions are shown as yellow spheres. The active site is indicated by a dashed box (left) and shown as a zoomed-in view (right). The likely rearrangements of key residues (D600, D602 and H603) that would need to occur for BLACT$_{Mg}$ to adopt an active conformation (coordinated by two $Mn^{2+}$ ions) are indicated by arrows. **b** Superposition of BLACT$_{Mg}$ (blue) and RNAse J from *S. coelicolor* (pink, PDB ID: 5A0T)[1], displayed as in panel **a**.

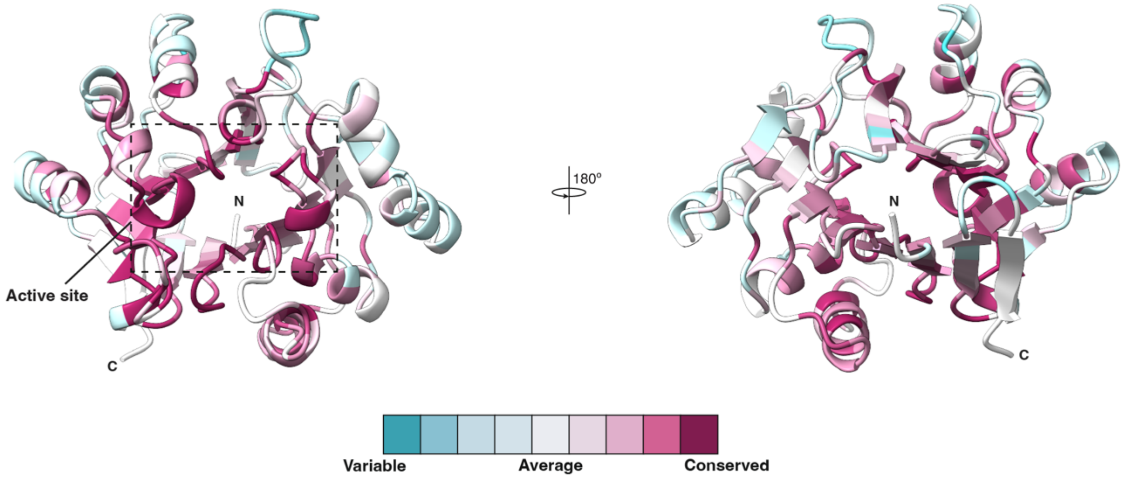

**Fig. S4: ConSurf analysis of BLACT$_{Mg}$**

Two orientations of BLACT$_{Mg}$ are shown in ribbon representation with residues coloured according to their conservation (cyan, low conservation; purple, high conservation). The active site of BLACT$_{Mg}$ is indicated by a dashed box (left) and is highly conserved. These images were generated using the ConSurf server[4].

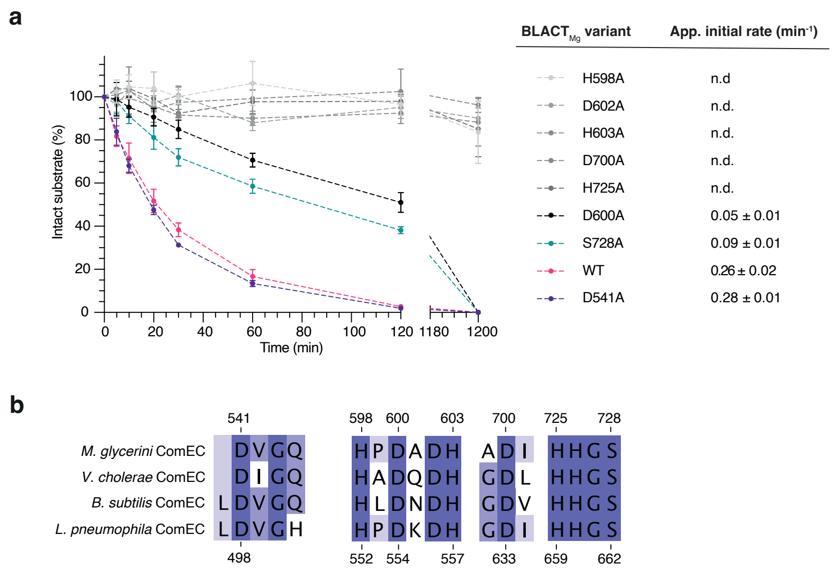

c

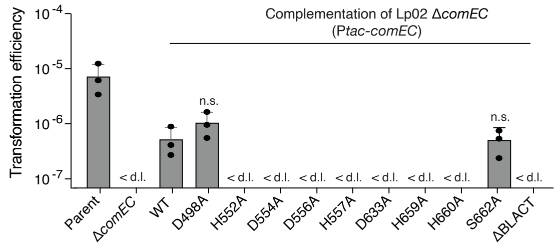

d

**Metal coordinating residues**

| BBL numbering | *M. glycerini* | *L. pneumophila* |
|---|---|---|
| H116 | H598 | H552 |
| H118 | D600 | D554 |
| D120 | D602 | D556 |
| H121 | H603 | H557 |
| D221 | D700 | D633 |
| H263 | H726 | H660 |

**Fig. S5: BLACT$_{Mg}$ nuclease activity is essential for natural transformation**
**a** Time-course nuclease activity assay monitoring the degradation of ssDNA by BLACT$_{Mg}$ variants. Nuclease assays were conducted with 1 μM wild-type BLACT$_{Mg}$ and 10 μM ssDNA substrates, 50 nucleotides (nt) in length. DNA substrates were fluorescently labelled with fluorescein (FAM) on the first base (thymine, T) at the 5' terminus. Error bars represent the standard error from three technical replicates. WT, wild type. **b** Multiple sequence alignment of the active site of BLACT$_{Mg}$ from *M. glycerini, Vibrio cholerae, B. subtilis and Legionella pneumophila*. Conservation is indicated by different shades of blue. **c** Transformation efficiencies of parental Lp02, Lp02 *ΔcomEC*, Lp02 *ΔBLACT* and Lp02 *ΔcomEC* strains complemented by ectopic expression of wild-type and mutant versions of ComEC corresponding to the mutations tested in panel **a**. Mean transformation efficiencies of three independent biological replicates are shown with error bars representing the standard deviation (SD). < d.l., below detection limit (d.l.). Statistical differences were determined on log-transformed data using an unpaired two-sided t-test with Welch's correction. The Lp02 strain complemented with wild-type ComEC was compared to those strains complemented with ComEC mutants. n.s., not statistically significant. **d** Table with metal coordinating residues according to the class B β-lactamases (BBL) numbering[5] and the corresponding residues in *M. glycerini* and *L. pneumophila*.

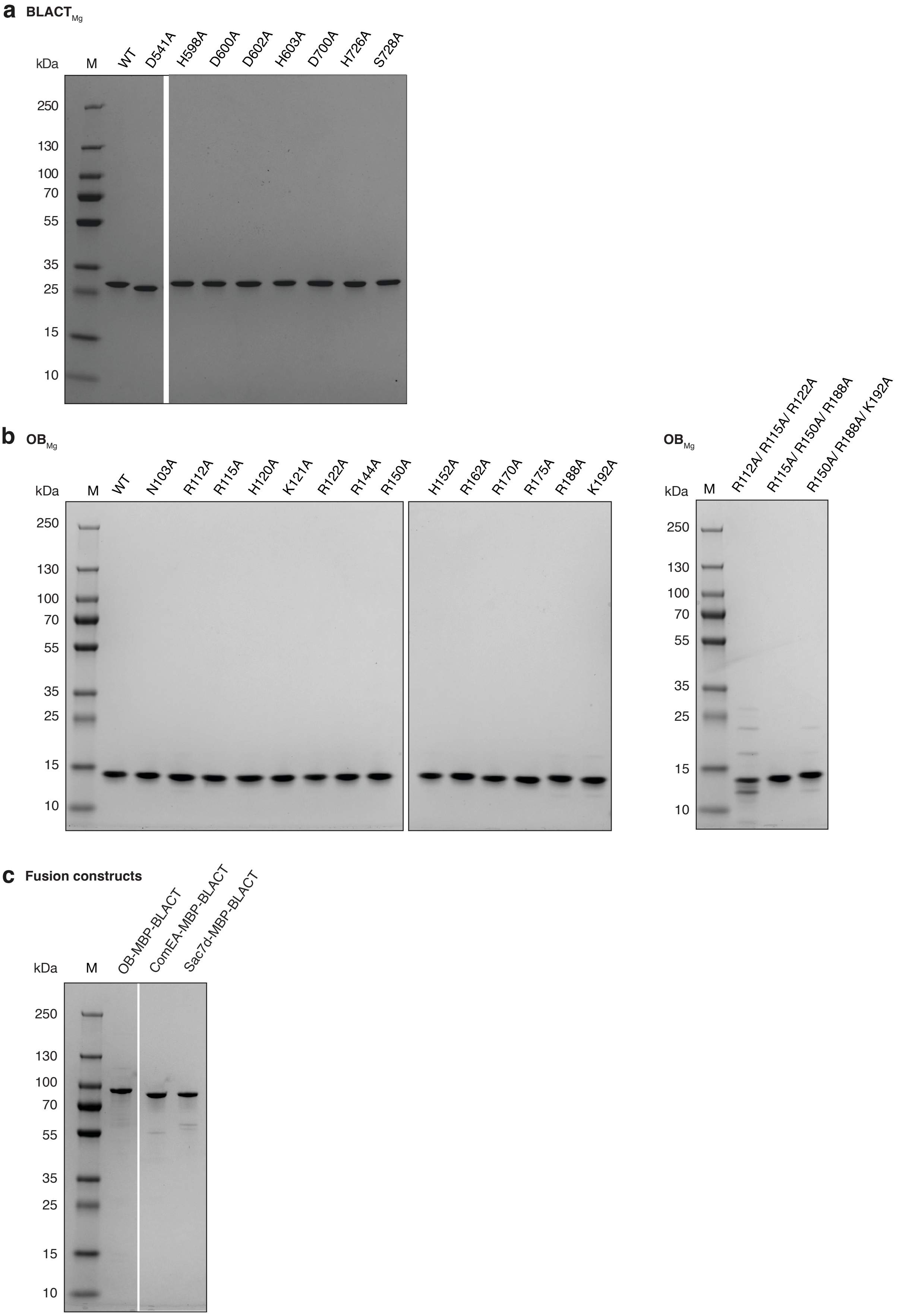
a
BLACT$_{Mg}$
kDa
M
WT
D541A
H598A
D600A
D602A
H603A
D700A
H726A
S728A
250
130
100
70
55
35
25
15
10
b
OB$_{Mg}$
kDa
M
WT
N103A
R112A
R115A
H120A
K121A
R122A
R144A
R150A
H152A
R162A
R170A
R175A
R188A
K192A
250
130
100
70
55
35
25
15
10
OB$_{Mg}$
kDa
M
R112A/ R115A/ R122A
R115A/ R150A/ R188A
R150A/ R188A/ K192A
250
130
100
70
55
35
25
15
10
c
Fusion constructs
kDa
M
OB-MBP-BLACT
ComEA-MBP-BLACT
Sac7d-MBP-BLACT
250
130
100
70
55
35
25
15
10

**Fig. S6: Purified proteins used in this study**

**a-c** Purified $BLACT_{Mg}$ (**a**), OB fold proteins from *M. glycerini* ($OB_{Mg}$) (**b**), and fusion constructs (**c**), resolved by SDS-PAGE. The OB, ComEA and BLACT domains within the fusion constructs are derived from *M. glycerini*, and Sac7d originates from *Sulfolobus acidocaldarius.* Further construct information can be found **Table S4**.

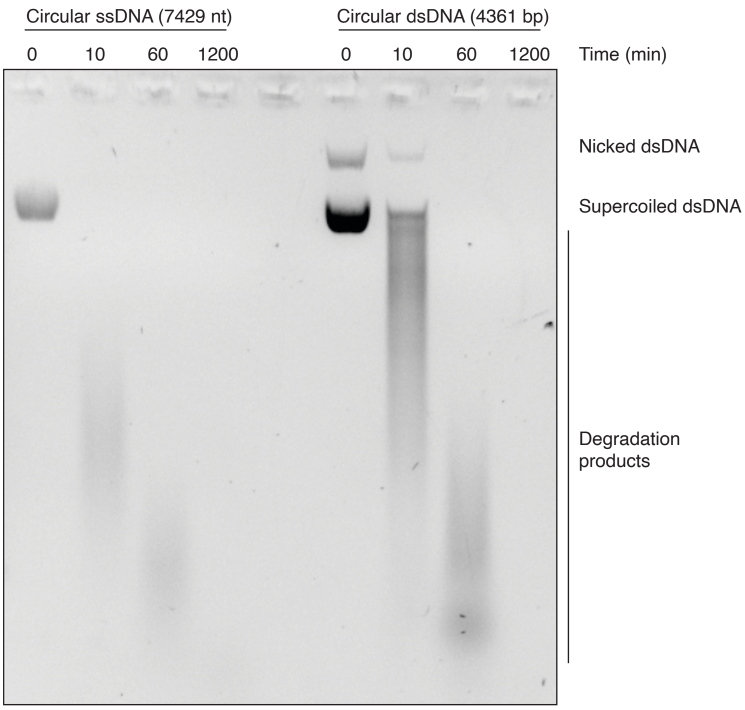

**Fig. S7: BLACT$_{Mg}$ degrades circular DNA *in vitro***
Time course nuclease assay resolved on a 1% (w/v) agarose gel showing the BLACT$_{Mg}$-dependent degradation of circular ssDNA and dsDNA *in vitro*. Substrate concentrations were adjusted such that the number of cleavable bonds were identical between ssDNA and dsDNA reactions. Different forms of DNA are labelled to the right of the gel.

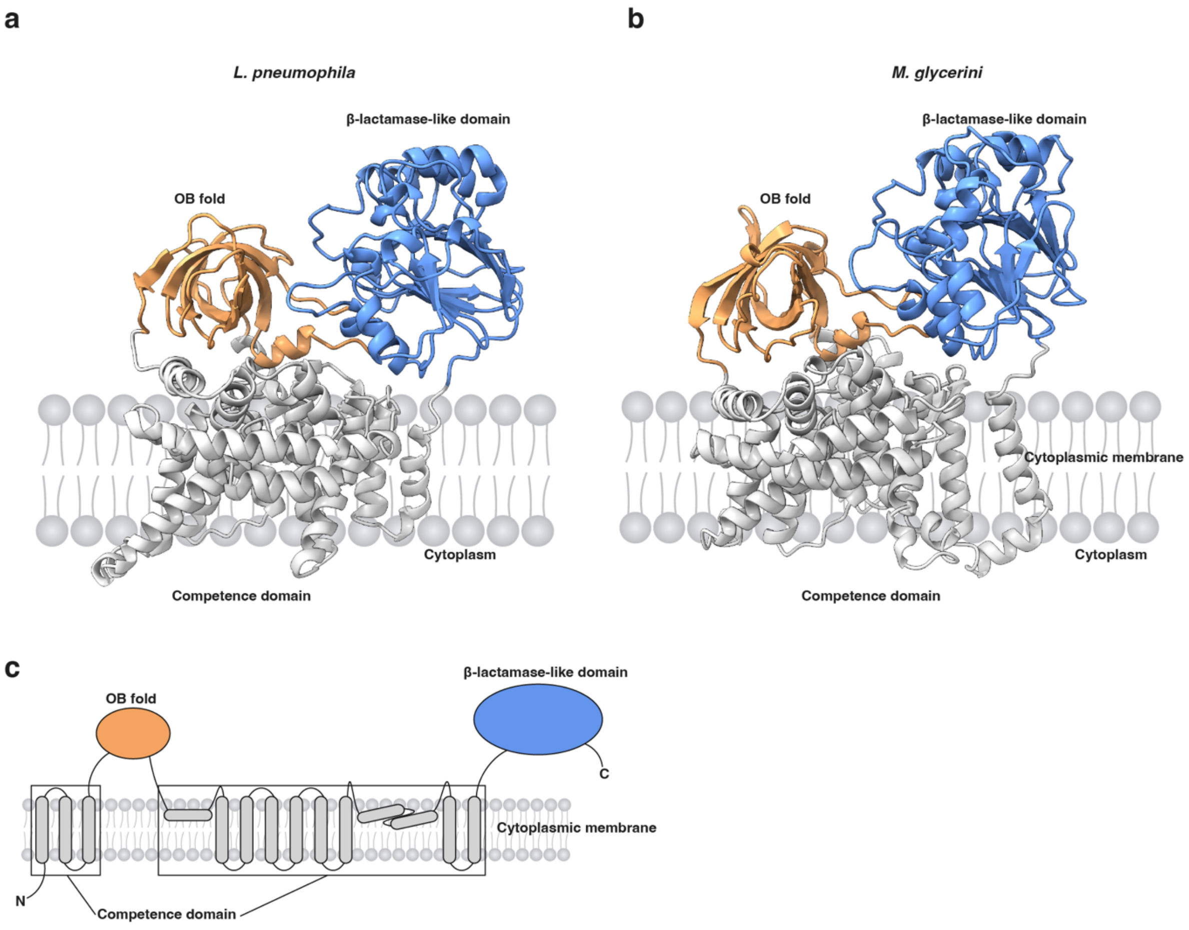

**Fig. S8: Predicted three-dimensional domain organisation of full-length ComEC**
**a, b** Structural predictions of *L. pneumophila* (**a**) and *M. glycerini* (**b**) ComEC orthologues using AlphaFold3[REF6] , shown in ribbon representation (OB fold, orange; β-lactamase-like domain, blue; competence domain, grey). **c** Schematic domain architecture of ComEC. The domains are coloured as in panels **a** and **b**.

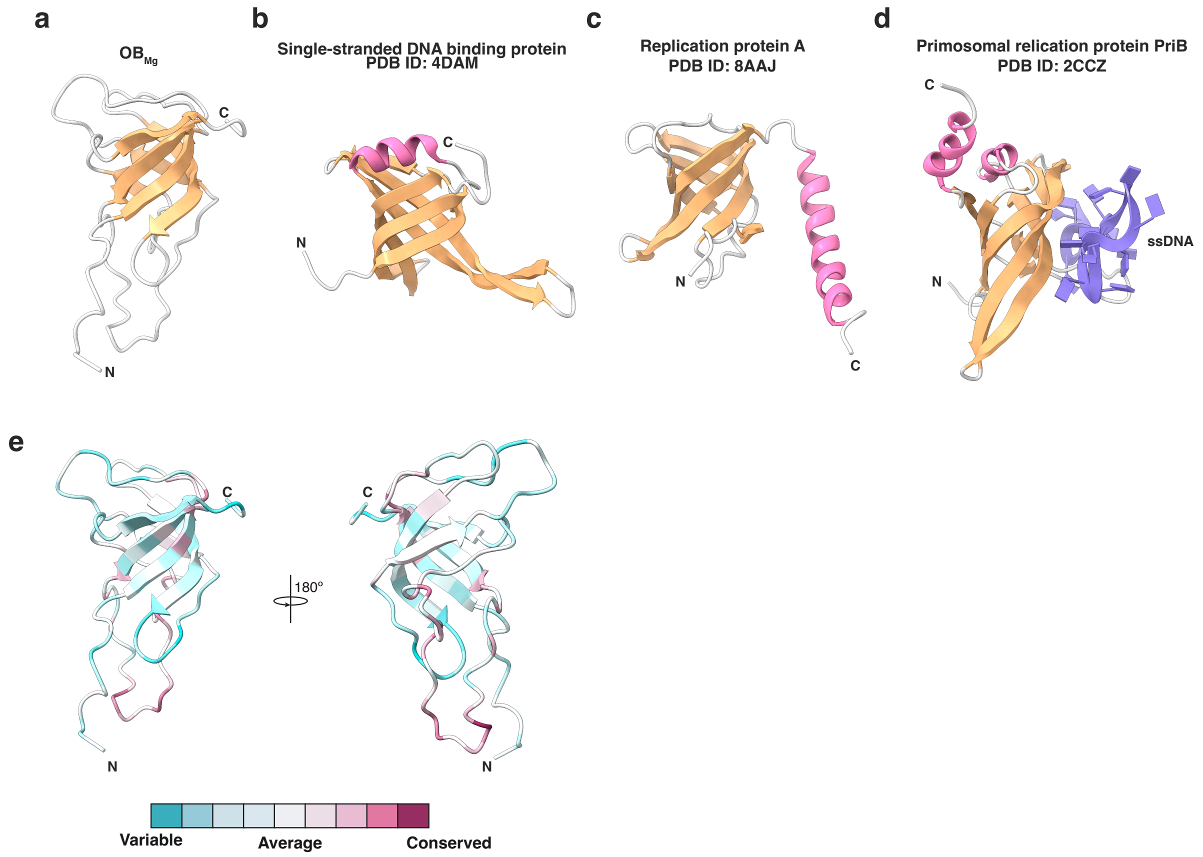

**Fig. S9: Structures of OB fold-containing proteins and ConSurf analysis of $OB_{Mg}$**

**a-d** Structural comparison of OB folds from different proteins, including $OB_{Mg}$ (this study) (**a**) single-stranded DNA binding protein from *S. coelicolor* (PDB ID: 4DAM)[7] (**b**), replication protein A from *Pyrococcus abyssi* (PDB ID: 8AAJ)[8] (**c**), and primosomal replication protein B (PriB) from *Escherichia coli* (PDB ID: 2CCZ)[9] (**d**). Structures are shown in ribbon representation (β-strands, orange; α-helices, pink; loops, grey; ssDNA, purple). **e** Two orientations of $OB_{Mg}$ with residues coloured according to conservation as in **Fig. S3**. Figure generated with the ConSurf server[4] .

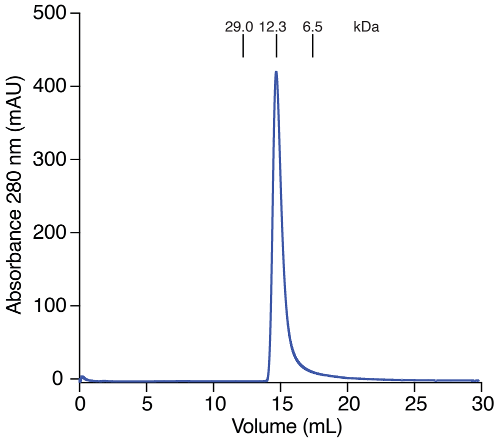

**Fig. S10: $OB_{Mg}$ is a monomer in solution**
Analytical size exclusion chromatography showing a single, monodisperse peak for $OB_{Mg}$ eluting at 14 mL. Molecular weights of globular proteins and their corresponding elution volumes, according to the manufacturer, are indicated above the chromatogram.

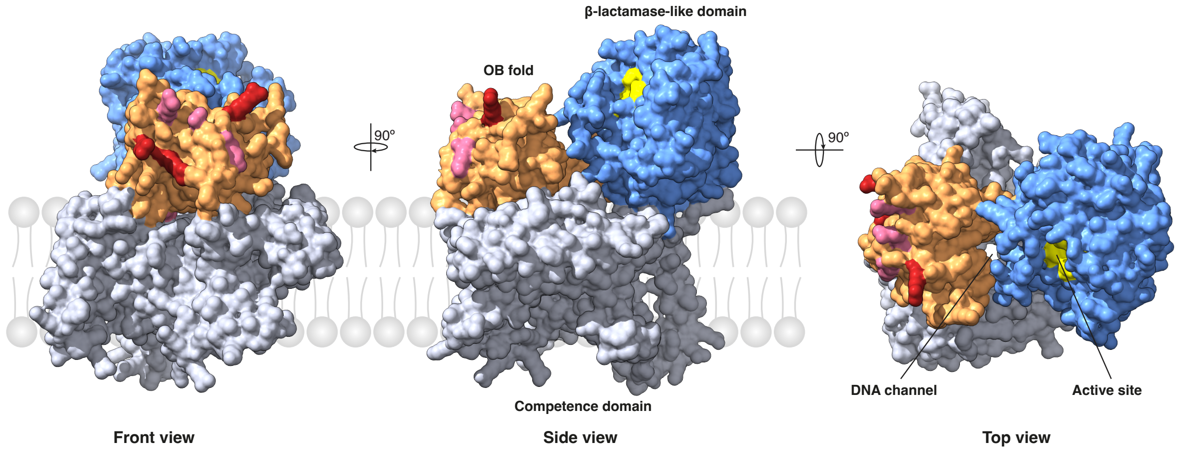

**Fig. S11: DNA binding residues are solvent exposed on the predicted structure of full-length ComEC**

The front, side and top view of full-length of ComEC from *M. glycerini* ($ComEC_{Mg}$) predicted by AlphaFold3[REF6] shown in surface representation (OB fold, orange; β-lactamase-like domain, blue; competence domain, grey). DNA binding residues identified in **Fig. 3d, e** coloured in shades of red. The active site of the β-lactamase-like domain and the DNA channel are also indicated on the figure.

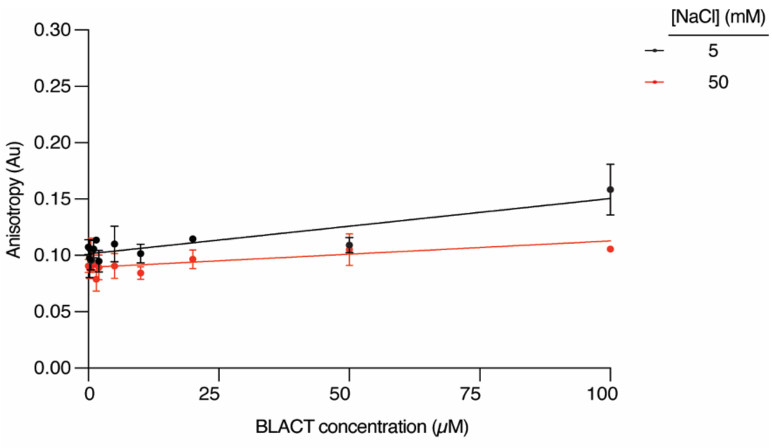

**Fig. S12: BLACT$_{Mg}$ does not bind DNA**
Fluorescence anisotropy binding experiment testing the interaction between 12 bp FAM-labelled dsDNA containing nuclease-resistant phosphorothioate (PTO) bonds and BLACT$_{Mg}$. The experiment was conducted in buffer containing 5 mM and 50 mM NaCl. Each experiment was performed in triplicate and error bars represent the standard error of the mean.

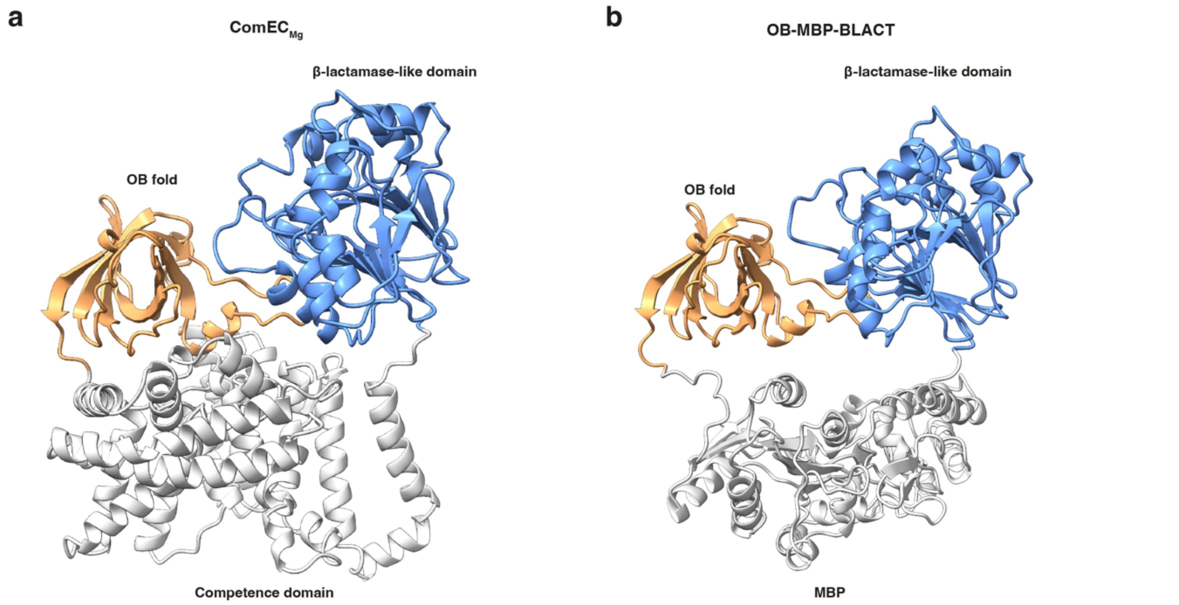

**Fig. S13: OB-MBP-BLACT displays a similar predicted domain organisation as ComEC**
**a, b** Structural prediction of ComEC$_{Mg}$ (**a**) and OB-MBP-BLACT (**b**) shown in ribbon representation (OB fold, orange; β-lactamase-like domain, blue; competence domain/MBP, grey).

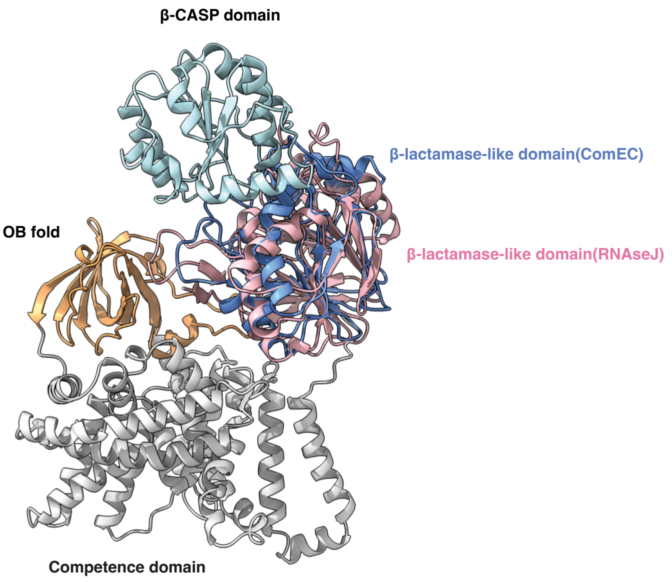

**Fig. S14: The relative positioning of the OB fold and β-CASP domains in ComEC and RNAse J**
Superposition of RNAse J from *S. coelicolor* (PDB ID: 5A0T)[1] and $ComEC_{Mg}$ using the β-lactamase-like domain to align the structures. Structures are displayed in ribbon representation (OB fold, orange; competence domain, grey; β-CASP domain, cyan; β-lactamase-like domain from $ComEC_{Mg}$, blue; β-lactamase-like domain from RNAse J, pink).

**Table S1: Data collection and refinement statistics of $BLACT_{Mg}$**

| **Protein** | $BLACT_{Mg}$ |
|---|---|
| **Data collection** | |
| Wavelength (Å) | 0.999989 |
| Space group | $C222_1$ |
| Cell dimensions | |
| *a, b, c* (Å) | 80.957, 96.849, 71.719 |
| $\alpha, \beta, \gamma$ (°) | 90, 90, 90 |
| Resolution (Å) | 62.11-1.86 (1.90-1.86) |
| $R_{sym}$ or $R_{merge}$ | 0.069(0.352) |
| Completeness (%) | 99.52(99.16) |
| Redundancy | 6.6(6.9) |
| Wilson B-factor | 22.46 |
| **Refinement** | |
| Resolution (Å) | 62.11-1.86 (1.90-1.86) |
| Unique reflections | 23812 |
| $R_{work}$ / $R_{free}$ | 0.177/0.212 |
| No. atoms (non-hydrogen) | |
| Protein | 2053 |
| Ligand | 5 |
| Water | 137 |
| Protein residues | 266 |
| Average B-factors | 29.99 |
| B-factors | |
| Protein | 29.72 |
| Ligand | 54.49 |
| Water | 33.16 |
| R.m.s deviations | |
| Bond lengths (Å) | 0.0056 |
| Bond angles (°) | 0.85 |
| Ramachandran plot | |
| Favored (%) | 95.08 |
| Allowed (%) | 4.92 |
| Outliers (%) | 0 |
| Rotamer outliers (%) | 0.46 |
| Clashscore | 1.71 |
| **PDB code** | XXXX |

Single crystal is used for the diffraction measurement.
Numbers in brackets are for the highest resolution shell.

**Table S2: NMR structural statistics of $OB_{Mg}$**

| **Protein** | $OB_{Mg}$ |
|---|---|
| **NMR distance and dihedral constraints** | |
| Distance constraints | |
| Total NOE | 2098 |
| Intra-residue | 379(18.1%) |
| Inter-residue | |
| Sequential ($\|i-j\| = 1$) | 887(42.3%) |
| Medium-range ($\|i-j\| < 5$) | 214(10.2%) |
| Long-range ($\|i-j\| \geq 5$) | 997(47.5%) |
| Hydrogen bonds | 0 |
| Total dihedral angle restraints | 140 |
| $\phi$ | 70 |
| $\psi$ | 70 |
| **Structure statistics** | |
| Violations (mean) | |
| Distance constraints (Å) | 2(0.20) |
| Dihedral angle constraints (°) | 4 (21) |
| Max. dihedral angle violation (°) | 22 |
| Max. distance constraint violation (Å) | 0.39 |
| Deviations from idealized geometry | |
| Bond lengths (Å) | 0.0036 ± 0.0012 |
| Bond angles (°) | 1.482 ± 0.494 |
| Average pairwise r.m.s. deviation** (Å) | |
| Heavy | 0.49 ± 0.11 |
| Backbone | 0.85 ± 0.14 |

* Statistics computed for the deposited bundle of 20 violation energy best structures selected out of 30 amber energy best structures.

** Pairwise r.m.s. deviation for the structured regionds comprising residues 11–80 and 107–124 was calculated among 20 out of 30 refined structures.

#### Supplementary Table 3: Bacterial strains used in this study

| Name | Relevant genotype/description | Source/Reference |
|---|---|---|
| *E. coli* | | |
| BL21 (DE3) | *E. coli* expression strain | NEB (cat. no. C2527H/I) |
| Stellar (HST08 strain) | *E. coli* cloning strain | Takara (cat. no. 636763/636766) |
| DH5α λpir | *E. coli cloning strain:* Encodes π protein for the replication of the *pir*-dependent origin of replication - *oriR*(*R6K)* | [10,11] |
| *L. pneumophila* | | |
| Lp02 WT | Philadelphia-1 *rpsL hsdR thyA*; SmR | [12] |
| Lp02 *ΔcomEC* | Lp02 Δ*comEC* | [13] |

**Supplementary Table 4: Plasmids used in this study**

| Name | Relevant genotype/description | Source/Reference |
|---|---|---|
| pMMB207C | *Legionella* expression vector derived from RSF1010: IncQ lacIq *Ptac oriT* Δ*mobA*; CmR | [14] |
| pMMB207C-*comEC$_{Lp}$* | *L. pneumophila* wild-type *comEC* | This study |
| pMMB207C-*comEC$_{Lp}$* D498A | pMMB207C-*comEC$_{Lp}$*, with *comEC* D498A mutation | This study |
| pMMB207C-*comEC$_{Lp}$* H552A | pMMB207C-*comEC$_{Lp}$*, with *comEC* H552A mutation | This study |
| pMMB207C-*comEC$_{Lp}$* D554A | pMMB207C-*comEC$_{Lp}$*, with *comEC* D554A mutation | This study |
| pMMB207C-*comEC$_{Lp}$* D556A | pMMB207C-*comEC$_{Lp}$*, with *comEC* D556A mutation | This study |
| pMMB207C-*comEC$_{Lp}$* H557A | pMMB207C-*comEC$_{Lp}$*, with *comEC* *H*557A mutation | This study |
| pMMB207C-*comEC$_{Lp}$* D663A | pMMB207C-*comEC$_{Lp}$*, with *comEC* D663A mutation | This study |
| pMMB207C-*comEC$_{Lp}$* H659A | pMMB207C-*comEC$_{Lp}$*, with *comEC* H659A mutation | This study |
| pMMB207C-*comEC$_{Lp}$* H660A | pMMB207C-*comEC$_{Lp}$*, with *comEC* H660A mutation | This study |
| pMMB207C-*comEC$_{Lp}$* S662A | pMMB207C-*comEC$_{Lp}$*, with *comEC* S662A mutation | This study |
| pMMB207C-*comEC$_{Lp}$* ΔBLACT | pMMB207C-*comEC$_{Lp}$*, with *comEC* BLACT deletion (Δ489-735) | This study |
| pMMB207C-*comEC$_{Lp}$* F523A | pMMB207C-*comEC$_{Lp}$*, with *comEC* F523A mutation | This study |
| pMMB207C-*comEC$_{Lp}$* Y522A/F523A | pMMB207C-*comEC$_{Lp}$*, with *comEC* Y522A/F523A mutation | This study |
| pMMB207C-*comEC$_{Lp}$* R688A | pMMB207C-*comEC$_{Lp}$*, with *comEC* R688A mutation | This study |
| pMMB207C-*comEC$_{Lp}$* F689A | pMMB207C-*comEC$_{Lp}$*, with *comEC* F689A mutation | This study |
| pMMB207C-*comEC$_{Lp}$* F691A | pMMB207C-*comEC$_{Lp}$*, with *comEC* F691A mutation | This study |
| pMMB207C-*comEC$_{Lp}$* F689A/F691A | pMMB207C-*comEC$_{Lp}$*, with *comEC* F689A/F691A mutation | This study |
| pMMB207C-*comEC$_{Lp}$* Y108A | pMMB207C-*comEC$_{Lp}$*, with *comEC* Y108A mutation | This study |
| pMMB207C-*comEC$_{Lp}$* Y140A | pMMB207C-*comEC$_{Lp}$*, with *comEC* Y140A mutation | This study |
| pMMB207C-*comEC$_{Lp}$* Y154A | pMMB207C-*comEC$_{Lp}$*, with *comEC* Y154A mutation | This study |
| pMMB207C-*comEC$_{Lp}$* W212A | pMMB207C-*comEC$_{Lp}$*, with *comEC* W212A mutation | This study |
| pMMB207C-*comEC$_{Lp}$* F331A | pMMB207C-*comEC$_{Lp}$*, with *comEC* F331A mutation | This study |
| pOPINS | *E. coli* expression vector, N-terminal $His_6$-SUMO tag, T7 promoter, KanR | [15] |
| pOPINS-*BLACT$_{Mg}$* | *M. glycerini* β-lactamase-like domain (532-797) | This study |
| pOPINS-*BLACT$_{Mg}$* D541A | pOPINS-*BLACT$_{Mg}$*, with BLACT D541A mutation | This study |
| pOPINS-*BLACT$_{Mg}$* H598A | pOPINS-*BLACT$_{Mg}$*, with BLACT H598A mutation | This study |
| pOPINS-*BLACT$_{Mg}$* D600A | pOPINS-*BLACT$_{Mg}$*, with BLACT D600A mutation | This study |

| | | |
|---|---|---|
| pOPINS-*BLACT*$_{Mg}$ D602A | pOPINS-*BLACT*$_{Mg}$, with BLACT D602A mutation | This study |
| pOPINS-*BLACT*$_{Mg}$ H603A | pOPINS-*BLACT*$_{Mg}$, with BLACT H603A mutation | This study |
| pOPINS-*BLACT*$_{Mg}$ D700A | pOPINS-*BLACT*$_{Mg}$, with BLACT D700A mutation | This study |
| pOPINS-BLACT$_{Mg}$ H725A | pOPINS-*BLACT*$_{Mg}$, with BLACT H725A mutation | This study |
| pOPINS-*BLACT*$_{Mg}$ S728A | pOPINS-*BLACT*$_{Mg}$, with BLACT S728A mutation | This study |
| pOPINS-*OB*$_{Mg}$ | *M. glycerini* OB fold (76-199) | This study |
| pOPINS-*OB*$_{Mg}$ N103A | pOPINS-*OB*$_{Mg}$, with N103A mutation | This study |
| pOPINS-*OB*$_{Mg}$ R112A | pOPINS-*OB*$_{Mg}$, with R112A mutation | This study |
| pOPINS-*OB*$_{Mg}$ R115A | pOPINS-*OB*$_{Mg}$, with R115A mutation | This study |
| pOPINS-*OB*$_{Mg}$ H120A | pOPINS-*OB*$_{Mg}$, with H120A mutation | This study |
| pOPINS-*OB*$_{Mg}$ K121A | pOPINS-*OB*$_{Mg}$, with K121A mutation | This study |
| pOPINS-*OB*$_{Mg}$ R122A | pOPINS-*OB*$_{Mg}$, with R122A mutation | This study |
| pOPINS-*OB*$_{Mg}$ R144A | pOPINS-*OB*$_{Mg}$, with R144A mutation | This study |
| pOPINS-*OB*$_{Mg}$ R150A | pOPINS-*OB*$_{Mg}$, with R150A mutation | This study |
| pOPINS-*OB*$_{Mg}$ H152A | pOPINS-*OB*$_{Mg}$, with H152A mutation | This study |
| pOPINS-*OB*$_{Mg}$ R162A | pOPINS-*OB*$_{Mg}$, with R162A mutation | This study |
| pOPINS-*OB*$_{Mg}$ R170A | pOPINS-*OB*$_{Mg}$, with R170A mutation | This study |
| pOPINS-*OB*$_{Mg}$ R175A | pOPINS-*OB*$_{Mg}$, with R175A mutation | This study |
| pOPINS-*OB*$_{Mg}$ R188A | pOPINS-*OB*$_{Mg}$, with R188A mutation | This study |
| pOPINS-*OB*$_{Mg}$ K192A | pOPINS-*OB*$_{Mg}$, with K192A mutation | This study |
| pOPINS-*OB*$_{Mg}$ R112A/R115A/R122A | pOPINS-*OB*$_{Mg}$, with R112A, R115A, R122A mutations | This study |
| pOPINS-*OB*$_{Mg}$ R115A/R150A/R188A | pOPINS-*OB*$_{Mg}$, with R115A, R150A, R188A mutations | This study |
| pOPINS-*OB*$_{Mg}$ R150A/R188A/K192A | pOPINS-*OB*$_{Mg}$, with R150A, R188A, K192A mutations | This study |
| pOPINS-*OB-MBP-BLACT* | OB-MBP-BLACT fusion protein (OB and BLACT domains from *M. glycerini*) | This study |
| pOPINS-*ComEA-MBP-BLACT* | ComEA-MBP-BLACT fusion protein (ComEA and BLACT domains from *M. glycerini*) | This study |
| pOPINS-*Sac7d-MBP-BLACT* | Sac7d-MBP-BLACT fusion protein (Sac7d from *S. acidocaldarius* and BLACT from *M. glycerini*) | This study |
| pTRC99A-*lpg2953-2958::Kan* | *L. pneumophila* genomic region spanning *lpg2953-2958*. The *hipB* gene (*lpg2955*) is disrupted by the kanamycin resistance gene, Kan | [13] |

**Supplementary Table 5: Oligonucleotides used in this study**

| Name | Sequence (5' to 3') | Construct |
|---|---|---|
| *Cloning* | | |
| Lpg2953-Fw | ATCTCTGGTGTGTTCGGATAGAT TATGCGAGAGGTCTATTTGAAGA TTCTCTGACTATG | pTRC99A-*lpg2953-2958::Kan* amplification of tDNA for transformation assay |
| Lpg2958-Rv | GTCGACTCTAGACACAGACATG GCCTGGAAACGTTGGTGGG | pTRC99A-*lpg2953-2958::Kan* amplification of tDNA for transformation assay |
| pMMB207C-*comEC$_{Lp}$*-ΔBLACT-Fw | CACCTCGTACTGCAATTTATTAA ATGGATTGGCTGACCCATGTTAT ATCTAAGCC | pMMB207C-*comEC$_{Lp}$*-ΔBLACT |
| pMMB207C-*comEC$_{Lp}$*-ΔBLACT-Rv | ATAAATTGCAGTACGAGGTGGA AATAAGGGAAGG | pMMB207C-*comEC$_{Lp}$*-ΔBLACT |
| pMMB207C-*comEC$_{Lp}$*-D498A-Fw | AATATTTTGGCTGTTGGGCAAGG ACTTG | pMMB207C-*comEC$_{Lp}$*-D498A |
| pMMB207C-*comEC$_{Lp}$*-D498A-Rv | AATATTTTGGCTGTTGGGCAAGG ACTTG | pMMB207C-*comEC$_{Lp}$*-D498A |
| pMMB207C-*comEC$_{Lp}$*-H552A-Fw | GTAATAAGCGCTCCCGATAAAG ATCATAAAGGCG | pMMB207C-*comEC$_{Lp}$*-H552A |
| pMMB207C-*comEC$_{Lp}$*-H552A-Rv | TTATCGGGAGCGCTTATTACTAT TTTGTCAATTGTTTTTATTCC | pMMB207C-*comEC$_{Lp}$*-H552A |
| pMMB207C-*comEC$_{Lp}$*-D554A-Fw | CATCCCGCTAAAGATCATAAAGG CGG | pMMB207C-*comEC$_{Lp}$*-D554A |
| pMMB207C-*comEC$_{Lp}$*-D554A-Rv | TATGATCTTTAGCGGGATGGCTT ATTACTATTTTGTCAATTG | pMMB207C-*comEC$_{Lp}$*-D554A |
| pMMB207C-*comEC$_{Lp}$*-D556A-Fw | CCGATAAAGCTCATAAAGGCGG ATTAAATTCCC | pMMB207C-*comEC$_{Lp}$*-D556A |
| pMMB207C-*comEC$_{Lp}$*-D556A-Rv | GCCTTTATGAGCTTTATCGGGAT GGCTTATTACTATTTTG | pMMB207C-*comEC$_{Lp}$*-D556A |
| pMMB207C-*comEC$_{Lp}$*-H557A-Fw | CCGATAAAGATGCTAAAGGCGG ATTAAATTCCC | pMMB207C-*comEC$_{Lp}$*-H557A |
| pMMB207C-*comEC$_{Lp}$*-H557A-Rv | GCCTTTAGCATCTTTATCGGGAT GGCTTATTACTATTTTG | pMMB207C-*comEC$_{Lp}$*-H557A |
| pMMB207C-*comEC$_{Lp}$*-D663A-Fw | AACGGGAGCTATTGAAAAAGCA GCTGAAGAGTAC | pMMB207C-*comEC$_{Lp}$*-D663A |
| pMMB207C-*comEC$_{Lp}$*-D663A-Rv | GCTTTTTCAATAGCTCCCGTTAA CAAAACCCTGC | pMMB207C-*comEC$_{Lp}$*-D663A |
| pMMB207C-*comEC$_{Lp}$*-H659A-Fw | GTGCCTGCTCATGGGAGCAAAA CGTC | pMMB207C-*comEC$_{Lp}$*-H659A |
| pMMB207C-*comEC$_{Lp}$*-H659A-Rv | GCTCCCATGAGCAGGCACAACT AAAA | pMMB207C-*comEC$_{Lp}$*-H659A |
| pMMB207C-*comEC$_{Lp}$*-H660A-Fw | GCCTCATGCTGGGAGCAAAACG TCTTC | pMMB207C-*comEC$_{Lp}$*-H660A |
| pMMB207C-*comEC$_{Lp}$*-H660A-Rv | GCTCCCAGCATGAGGCACAACT AAAATTTCGGATGCCAG | pMMB207C-*comEC$_{Lp}$*-H660A |
| pMMB207C-*comEC$_{Lp}$*-S662A-Fw | CATCATGGGGCTAAAACGTCTTC ATCC | pMMB207C-*comEC$_{Lp}$*-S662A |
| pMMB207C-*comEC$_{Lp}$*-S662A-Rv | GACGTTTTAGCCCCATGATGAG GCAC | pMMB207C-*comEC$_{Lp}$*-S662A |
| pMMB207C-*comEC$_{Lp}$*-F523A-Fw | TTCCTACGCGCAGGGGAGTGAT TTAGGGCAAATGGC | pMMB207C-*comEC$_{Lp}$*-F523A |
| pMMB207C-*comEC$_{Lp}$*-F523A-Rv | TCCCCTGCGCGTAGGAATCTCC CGTGTCGTAGAGTAAAAC | pMMB207C-*comEC$_{Lp}$*-F523A |
| pMMB207C-*comEC$_{Lp}$*-Y522A/F523A-Fw | AGATTCCGCGGCGCAGGGGAGT GATTTAGGGCAAATGGC | pMMB207C-*comEC$_{Lp}$*-Y522A/F523A |

| | | |
|---|---|---|
| pMMB207C-*comEC$_{Lp}$*-Y522A/F523A-Rv | TCCCCTGCGCCGCGGAATCTCCCGTGTCGTAGAGTAAAACATG | pMMB207C-*comEC$_{Lp}$*-Y522A/F523A |
| pMMB207C-*comEC$_{Lp}$*-R688A-Fw | TGATAATGCGTTCAAATTTCCACATTCTAAAACTTTGCAAAGCATG | pMMB207C-*comEC$_{Lp}$*-R688A |
| pMMB207C-*comEC$_{Lp}$*-R688A-Rv | ATTTGAACGCATTATCAAATCCTAAGGATGCAATGGCATAAAGC | pMMB207C-*comEC$_{Lp}$*-R688A |
| pMMB207C-*comEC$_{Lp}$*-F689A-Fw | TAATCGCGCGAAATTTCCACATTCTAAAACTTTGCAAAGCATGAAAAC | pMMB207C-*comEC$_{Lp}$*-F689A |
| pMMB207C-*comEC$_{Lp}$*-F689A-Rv | GAAATTTCGCGCGATTATCAAATCCTAAGGATGCAATGGC | pMMB207C-*comEC$_{Lp}$*-F689A |
| pMMB207C-*comEC$_{Lp}$*-F691A-Fw | CTTCAAAGCGCCACATTCTAAAACTTTGCAAAGCATGAAAACATTAG | pMMB207C-*comEC$_{Lp}$*-F691A |
| pMMB207C-*comEC$_{Lp}$*-F691A-Rv | AATGTGGCGCTTTGAAGCGATTATCAAATCCTAAGGATGCAATG | pMMB207C-*comEC$_{Lp}$*-F691A |
| pMMB207C-*comEC$_{Lp}$*-F689A/F691A-Fw | TTGATAATCGCGCGAAAGCGCCACATTCTAAAACTTTGCAAAGCATGAAAACATTAG | pMMB207C-*comEC$_{Lp}$*-F689A/F691A |
| pMMB207C-*comEC$_{Lp}$*-F689A/F691A-Rv | TTAGAATGTGGCGCTTTCGCGCGATTATCAAATCCTAAGGATGCAATGGC | pMMB207C-*comEC$_{Lp}$*-F689A/F691A |
| pMMB207C-*comEC$_{Lp}$*-Y108A-Fw | AAACTGGGCGAACAAACCACCTGTCTTACGAGCC | pMMB207C-*comEC$_{Lp}$*-Y108A |
| pMMB207C-*comEC$_{Lp}$*-Y108A-Rv | GTTTGTTCGCCCAGTTTAATTGGATCAAGCCTTGAGCAG | pMMB207C-*comEC$_{Lp}$*-Y108A |
| pMMB207C-*comEC$_{Lp}$*-Y140A-Fw | TTTTAATGCGGTACGTTACCTGGCGGCTCG | pMMB207C-*comEC$_{Lp}$*-Y140A |
| pMMB207C-*comEC$_{Lp}$*-Y140A-Rv | AACGTACCGCATTAAAACCTCCCGGGTTGTGGAAG | pMMB207C-*comEC$_{Lp}$*-Y140A |
| pMMB207C-*comEC$_{Lp}$*-Y154A-Fw | GACTGGCGCGATTCGTTATCGGAATAATAAGCTAATTAGCAATCAACC | pMMB207C-*comEC$_{Lp}$*-Y154A |
| pMMB207C-*comEC$_{Lp}$*-Y154A-Rv | AACGAATCGCGCCAGTCCAATGGATATGACGAGC | pMMB207C-*comEC$_{Lp}$*-Y154A |
| pMMB207C-*comEC$_{Lp}$*-W212A-Fw | AGATCATGCGAATTTGTTCCGAAGAACAGGAACGATAC | pMMB207C-*comEC$_{Lp}$*-W212A |
| pMMB207C-*comEC$_{Lp}$*-W212A-Rv | ACAAATTCGCATGATCTGAACTGATATGGTGAGTAACATTTAACGTC | pMMB207C-*comEC$_{Lp}$*-W212A |
| pMMB207C-*comEC$_{Lp}$*-F331A-Fw | TTTTTACGCGTCATTTCTTGCCGTGGCCTGTC | pMMB207C-*comEC$_{Lp}$*-F331A |
| pMMB207C-*comEC$_{Lp}$*-F331A-Rv | GAAATGACGCGTAAAAACCTTGCAGGAATACGGCATG | pMMB207C-*comEC$_{Lp}$*-F331A |
| pOPINS-linearise-Fw | TAAAGCTTTCTAGACCATTTAAACACCACCACCACC | pOPINS fragment |
| pOPINS-linearise-Rv | ACCACCGATCTGTTCGCGATGCG | pOPINS fragment |
| pOPINS-*BLACT$_{Mg}$*-Fw | GCGAACAGATCGGTGGTCAAGAAGAGCTGAAAGTCACTTTCATCGATGTTGG | pOPINS-*BLACT$_{Mg}$* |
| pOPINS-*BLACT$_{Mg}$*-Rv | ATGGTCTAGAAAGCTTTAAATCGTTGTGTTGACTTGCCAACGCTC | pOPINS-*BLACT$_{Mg}$* |
| pOPINS-*BLACT$_{Mg}$*-D541A-Fw | GCGAACAGATCGGTGGTCAAGAAGAGCTGAAAGTCACTTTCATCGCGGTTGG | pOPINS-*BLACT$_{Mg}$* D541A |
| pOPINS-*BLACT$_{Mg}$*-D541A-Rv | ATGGTCTAGAAAGCTTTAAATCGTTGTGTTGACTTGCCAACGCTC | pOPINS-*BLACT$_{Mg}$* D541A |

| | | |
|---|---|---|
| pOPINS-$BLACT_{Mg}$-H598A-Fw | AGCACGGCGCCGGACGCGGACCACATCGG | pOPINS-$BLACT_{Mg}$ H598A |
| pOPINS-$BLACT_{Mg}$-H598A-Rv | GTCCGGCGCCGTGCTAACAACAACGTCTAAATGGCG | pOPINS-$BLACT_{Mg}$ H598A |
| pOPINS-$BLACT_{Mg}$-D600A-Fw | GTTAGCACGCATCCGGCGGCGGACCACATC | pOPINS-$BLACT_{Mg}$ D600A |
| pOPINS-$BLACT_{Mg}$-D600A-Rv | CTGCCAGACCACCGATGTGGTCCGCCGC | pOPINS-$BLACT_{Mg}$ D600A |
| pOPINS-$BLACT_{Mg}$-D602A-Fw | GACGCGGCGCACATCGGTGGTCTGGCAGCG | pOPINS-$BLACT_{Mg}$ D602A |
| pOPINS-$BLACT_{Mg}$-D602A-Rv | GATGTGCGCCGCGTCCGGATGCGTGCTAAC | pOPINS-$BLACT_{Mg}$ D602A |
| pOPINS-$BLACT_{Mg}$-H603A-Fw | GCGGACGCGATCGGTGGTCTGGCAGCGGTTG | pOPINS-$BLACT_{Mg}$ D603A |
| pOPINS-$BLACT_{Mg}$-H603A-Rv | ACCGATCGCGTCCGCGTCCGGATGCGTGC | pOPINS-$BLACT_{Mg}$ D603A |
| pOPINS-$BLACT_{Mg}$-D700A-Fw | GTCTGCTGTTGTCCGCTGCGATTGAAGCTGAG | pOPINS-$BLACT_{Mg}$ D700A |
| pOPINS-$BLACT_{Mg}$-D700A-Rv | CTTTTAAGTCTGCCATTGCCTCAGCTTCAATCGC | pOPINS-$BLACT_{Mg}$ D700A |
| pOPINS-$BLACT_{Mg}$-H725A-Fw | GCGTTCGACTGTATTTAAGGTACCAGCGCATGG | pOPINS-$BLACT_{Mg}$ H725A |
| pOPINS-$BLACT_{Mg}$-H725A-Rv | CAAGACCATAACGGGACCCATGCGCTGGTACC | pOPINS-$BLACT_{Mg}$ H725A |
| pOPINS-$BLACT_{Mg}$-S728A-Fw | GTATTTAAGGTACCACATCATGGGGCGCGTTATGGTC | pOPINS-$BLACT_{Mg}$ S728A |
| pOPINS-$BLACT_{Mg}$-S728A-Rv | GAGGAATTCACGCTCAAGACCATAACGCGCCC | pOPINS-$BLACT_{Mg}$ S728A |
| pOPINS-$OB_{Mg}$-Fw | GAACAGATCGGTGGTCATCAATCACGTCTCACGGGTGATCG | pOPINS-$OB_{Mg}$ |
| pOPINS-$OB_{Mg}$-Rv | GTCTAGAAAGCTTTAGTGACCTGGTTCGGTACCCAATTTCAC | pOPINS-$OB_{Mg}$ |
| pOPINS-$OB_{Mg}$-N103A-Fw | TATCCGGCGCGTGTGGTTTACACACTTGCCGCGCG | pOPINS-$OB_{Mg}$ N103A |
| pOPINS-$OB_{Mg}$-N103A-Rv | CACACGCGCCGGATACACACGAGGCTCTTCGATCAC | pOPINS-$OB_{Mg}$ N103A |
| pOPINS-$\mathrm{OB}_{\mathrm{Mg}}$-R112A-Fw | GCCGCGGCGGAAATTCGTCAAGGCGATTATCACAAGCG | pOPINS-$OB_{Mg}$ R112A |
| pOPINS-$OB_{Mg}$-R112A-Rv | AATTTCCGCCGCGGCAAGTGTGTAAACCACACGATTCGG | pOPINS-$OB_{Mg}$ R112A |
| pOPINS-$OB_{Mg}$-R115A-Fw | GAAATTGCGCAAGGCGATTATCACAAGCGTGTGCGCG | pOPINS-$OB_{Mg}$ R115A |
| pOPINS-$OB_{Mg}$-R115A-Rv | GCCTTGCGCAATTTCGCGCGCGGCAAGTGTGTAAA | pOPINS-$OB_{Mg}$ R115A |
| pOPINS-$OB_{Mg}$-H120A-Fw | GATTATGCGAAGCGTGTGCGCGAGAAAGTGCAAGTC | pOPINS-$OB_{Mg}$ H120A |
| pOPINS-$OB_{Mg}$-H120A-Rv | ACGCTTCGCATAATCGCCTTGACGAATTTCGCGCG | pOPINS-$OB_{Mg}$ H120A |
| pOPINS-$OB_{Mg}$-K121A-Fw | TATCACGCGCGTGTGCGCGAGAAAGTGCAAGTCGTAC | pOPINS-$OB_{Mg}$ K121A |
| pOPINS-$OB_{Mg}$-K121A-Rv | CACACGCGCGTGATAATCGCCTTGACGAATTTCGCGC | pOPINS-$OB_{Mg}$ K121A |
| pOPINS-$OB_{Mg}$-R122A-Fw | CACAAGGCGGTGCGCGAGAAAGTGCAAGTCGTACTGTATCGC | pOPINS-$OB_{Mg}$ R122A |
| pOPINS-$OB_{Mg}$-R122A-Rv | GCGCACCGCCTTGTGATAATCGCCTTGACGAATTTCGCGC | pOPINS-$OB_{Mg}$ R122A |
| pOPINS-$OB_{Mg}$-R144A-Fw | CTGTATGCGAGCGCGAAAGGTGGCGAGCCCGTGC | pOPINS-$OB_{Mg}$ R144A |

| | | |
|---|---|---|
| pOPINS-$OB_{Mg}$-R144A-Rv | CGCGCTCGCATACAGTACGACT TGCACTTTCTCGCGC | pOPINS-$OB_{Mg}$ R144A |
| pOPINS-$OB_{Mg}$-R150A-Fw | GTACTGGCGGTGCATGGTCAAC TTGCCGCACCCC | pOPINS-$OB_{Mg}$ R150A |
| pOPINS-$OB_{Mg}$-R150A-Rv | ATGCACCGCCAGTACATCACCA TAACGATAGAGCACGGG | pOPINS-$OB_{Mg}$ R150A |
| pOPINS-$OB_{Mg}$-H152A-Fw | CGTGTGGCGGGTCAACTTGCCG CACCCCCAGCTG | pOPINS-$OB_{Mg}$ H152A |
| pOPINS-$OB_{Mg}$-H152A-Rv | TTGACCCGCCACACGCAGTACA TCACCATAACGATAGAGCACGG | pOPINS-$OB_{Mg}$ H152A |
| pOPINS-$OB_{Mg}$-R162A-Fw | GCTGCGGCGAACCCCGGGGAG CTGGATTATCGTGCG | pOPINS-$OB_{Mg}$ H162A |
| pOPINS-$OB_{Mg}$-R162A-Rv | GGGGTTCGCCGCAGCTGGGGG TGCGGCAAGTTGAC | pOPINS-$OB_{Mg}$ H162A |
| pOPINS-$OB_{Mg}$-R170A-Fw | GATTATGCGGCGTATCTTGCAC GTCAATACATTTACAATCGC | pOPINS-$OB_{Mg}$ R170A |
| pOPINS-$OB_{Mg}$-R170A-Rv | ATACGCCGCATAATCCAGCTCC CCGGGGTTGCG | pOPINS-$OB_{Mg}$ R170A |
| pOPINS-$OB_{Mg}$-R175A-Fw | CTTGCAGCGCAATACATTTACAA TCGCATGCTTATTGACAATCCC | pOPINS-$OB_{Mg}$ R175A |
| pOPINS-$OB_{Mg}$-R175A-Rv | GTATTGCGCTGCAAGATACGCA CGATAATCCAGCTCCC | pOPINS-$OB_{Mg}$ R175A |
| pOPINS-$OB_{Mg}$-R188A-Fw | AATCCCGCGGCTATCGTGAAATT GGGTACCGAACC | pOPINS-$OB_{Mg}$ R188A |
| pOPINS-$OB_{Mg}$-R188A-Rv | GATAGCCGCGGGATTGTCAATA AGCATGCGATTGTAAATGTATTG ACG | pOPINS-$OB_{Mg}$ R188A |
| pOPINS-$OB_{Mg}$-K192A-Fw | ATCGTGGCGTTGGGTACCGAAC CAGGTCACTAAAGCTTTC | pOPINS-$OB_{Mg}$ R192A |
| pOPINS-$OB_{Mg}$-K192A-Rv | ACCCAACGCCACGATAGCACGG GGATTGTCAATAAGCATG | pOPINS-$OB_{Mg}$ R192A |
| pOPINS-$OB_{Mg}$-R112A/R115A/R122A-Fw | GCCGCGGCGGAAATTGCGCAAG GCGATTATCACAAG | pOPINS-$OB_{Mg}$ R112A/R115A/R122A |
| pOPINS-$OB_{Mg}$-R112A/R115A/R122A-Rv | AATTTCCGCCGCGGCAAGTGTG TAAACCACACGATTCGG | pOPINS-$OB_{Mg}$ R112A/R115A/R122A |
| pOPINS-$OB_{Mg}$-R150A/R188A/K192A-Fw | AATCCCGCGGCTATCGTGGCGT TGGGTACCGAACC | pOPINS-$OB_{Mg}$ R150A/R188A/K192A |
| pOPINS-$OB_{Mg}$-R150A/R188A/K192A-Rv | GATAGCCGCGGGATTGTCAATA AGCATGCGATTGTAAATGTATTG ACG | pOPINS-$OB_{Mg}$ R150A/R188A/K192A |
| pOPINS-$OB_{Mg}$-linearise-Fw | TAAAGCTTTCTAGACCATTTAAA CACCACCACCACC | pOPINS-$OB_{Mg}$ fragment |
| pOPINS-$OB_{Mg}$-linearise-Rv | GTGACCTGGTTCGGTACCCAAT TTCACG | pOPINS-$OB_{Mg}$ fragment |
| pOPINS-*OB-BLACT*-Fw | ACCGAACCAGGTCACCAAGAAG AGCTGAAAGTCACTTTCATCGAT G | pOPINS-*OB-BLACT* |
| pOPINS-*OB-BLACT*-Rv | GTCTAGAAAGCTTTAAATCGTTG TGTTGACTTGCCAACGC | pOPINS-*OB-BLACT* |
| pOPINS-*OB-BLACT*-linearise-Fw | GCGAACAGATCGGTGGTCAAGA AGAGCTGAAAGTCACTTTCATCG ATGTTGG | pOPINS-*OB-BLACT* fragment |
| pOPINS-*OB-BLACT*-linearise-Rv | GTGACCTGGTTCGGTACCCAAT TTCACG | pOPINS-*OB-BLACT* fragment |
| pOPINS-*OB-MBP-BLACT*-Fw | ACCGAACCAGGTCACGGCAGCA GCGGCAGCAGCAAAATCGAAGA AGGTAAACTGGTAATCTGG | pOPINS-*OB-MBP-BLACT* |

| | | |
|---|---|---|
| pOPINS-*OB-MBP-BLACT*-Rv | TTTCAGCTCTTCTTGGCTGCTGC<br>CGCTGCTGCCTTTCGTGATGCG<br>AGTCTGCGCGTCTTTCAGGGC | pOPINS-*OB-MBP-BLACT* |
| pOPINS-*MBP-BLACT*-linearise-Fw | GCAAAATCGAAGAAGGTAAACT<br>GGTAATCTGG | pOPINS-*MBP-BLACT* fragment |
| pOPINS-*MBP-BLACT*-linearise-Rv | ACCACCGATCTGTTCGCGATGC<br>GCTTCAATAA | pOPINS-*MBP-BLACT* fragment |
| pOPINS-*ComEA-MBP-BLACT*-Fw | ATCGCGAACAGATCGGTGGTGC<br>TAGAAAGATTAACATCAATACC | pOPINS-*ComEA-MBP-BLACT* |
| pOPINS-*ComEA-MBP-BLACT*-Rv | TTTACCTTCTTCGATTTTGCTGC<br>TGCCGCTGCTGCCGCGAACAGT<br>AATCAGATCTTTCAA | pOPINS-*ComEA-MBP-BLACT* |
| pOPINS-*Sac7d-MBP-BLACT*-Fw | ATCGCGAACAGATCGGTGGTAT<br>GGTGAAGGTGAAATTTAAATATA<br>AGGGA | pOPINS-*Sac7d-MBP-BLACT* |
| pOPINS-*Sac7d-MBP-BLACT*-Rv | TTTACCTTCTTCGATTTTGCTGC<br>TGCCGCTGCTGCCCTTTTTTTCA<br>CGTTCCGCGCGGGC | pOPINS-*Sac7d-MBP-BLACT* |
| *Biophysical assays*# | | |
| 5'OH-5'FAM* | T*GGGTAGGATCATCAGTAATAA<br>GGATAGTGGGAAAGCTCACAGA<br>CCACCT | Nuclease assay |
| 5'Phos-5'FAM* | pT*GGGTAGGATCATCAGTAATAA<br>GGATAGTGGGAAAGCTCACAGA<br>CCACCT | Nuclease assay |
| 5'OH-5'FAM*-(PTO) | T*/G/G/G/T/A/G/G/A/T/C/A/T/C/A/G<br>/T/A/A/T/A/A/G/G/A/T/A/G/T/G/G/G/<br>A/A/A/G/C/T/C/A/C/A/G/A/C/C/A/C/<br>C/G | Nuclease assay |
| 5'Phos-5'FAM*-(PTO) | pT*/G/G/G/T/A/G/G/A/T/C/A/T/C/A/<br>G/T/A/A/T/A/A/G/G/A/T/A/G/T/G/G/<br>G/A/A/A/G/C/T/C/A/C/A/G/A/C/C/A/<br>C/C/G | Nuclease assay |
| 5'OH-3'FAM£-(PTO) | C/G/G/G/T/A/G/G/A/T/C/A/T/C/A/G/<br>T/A/A/T/A/A/G/G/A/T/A/G/T/G/G/G/<br>A/A/A/G/C/T/C/A/C/A/G/A/C/C/A/C/<br>C/T£ | Nuclease assay |
| 5'OH-5'FAM*-(PTO) | T*/G/G/G/T/A/G/G/A/T/C/A/T/C/A/G<br>/T/A/A/T/A/A/G/G/A/T/A/G/T/G/G/G/<br>A/A/A/G/C/T/C/A/C/A/G/A/C/C/A/C/<br>C/G | Nuclease assay |
| 5'FAM$ | $GTTCGCAACGAA | Fluorescence anisotropy |
| 5'FAM$-(PTO) | $G/T/T/C/G/C/A/A/C/G/A/A | Fluorescence anisotropy |

### The complementary strand for dsDNA probes is not shown. Only one of the two strands is FAM-labelled. Unless stated otherwise, all DNA probes for nuclease assays are 50 nt in length, and all probes for fluorescence anisotropy are 12 nt in length.
* FAM was attached to the first base (thymine, T) at the 5′ terminus.
£ FAM was attached to the last base (thymine, T) at the 3′ terminus.
$ FAM was attached to the 5' phosphate group.
p 5' phosphorylation modification.
/ The uncleavable phosphorothioate bond.
/ The phosphodiester bond.
